# SYNPO2L Isoforms Regulate the Action Potential Characteristics and Contractility of Atrial Cardiomyocytes via YAP Signaling to Modulate Atrial Fibrillation Risk

**DOI:** 10.1101/2025.08.03.668361

**Authors:** Rishikesan Chandrakumar, Reva Shenwai, Farshad Farshidfar, Luke Zhang, Sylwia M. Figarska, Ana Budan, Olga Cisne-Thompson, Anley E. Tefera, Timothy Hoey, James R. Priest, Jonathan H. Tsui

**Author notes:** **Corresponding Author:** Jonathan Tsui, 171 Oyster Point Blvd., Suite 500, South San Francisco, CA 94080.

## Abstract

**Background:** SYNPO2L is a component of the sarcomere Z-disk with two distinct isoforms (SYNPO2L_A and SYNPO2L_B) described to regulate actin bundle size. Common and rare variations in SYNPO2L have been implicated in risk for heart failure and atrial fibrillation (AF); however, little is known about the function or pathophysiological mechanism upon disease risk.

**Methods:** The role of SYNPO2L was explored with human induced pluripotent stem cell cardiomyocyte (hiPSC-CM) models of a rare splicing mutation rs766868752 differentiated into both atrial and ventricular lineages. Electrophysiology was assessed via multi-electrode arrays (MEAs) and contractility was assessed using engineered heart tissues (EHTs). Genetic survival analyses were performed to explore the potential therapeutic role of the SYNPO2L_A isoform in AF risk after myocardial infarction.

**Results:** hiPSC-CMs carrying the splice mutation rs766868752 resulted in preferential expression of the SYNPO2L_B isoform and absence of SYNPO2L_A isoform, and atrial hiPSC-CMs displayed significant differences in action potential durations when unpaced, and spontaneous extra-systolic beats when paced. Contractility of atrial hiPSC-CMs carrying the splice mutation was severely compromised. Electrophysiological differences were normalized and contractility was partially restored by overexpression with AAV:SYNPO2L_A. Absence of SYNPO2L_A in the mutant led to an observed decreased phosphorylation of YAP along with significant downstream transcriptional effects. A direct interaction between SYNPO2L_A and LATS2, a known regulator of YAP phosphorylation was found to occur in hiPSC-CMs. Additionally, a common variant time-to-event analysis may suggest a beneficial effect of the SYNPO2L_A isoform to lower the risk of atrial fibrillation after myocardial infarction.

**Conclusions:** A rare splice mutation conferring isoform predominance of SYNPO2L_B appears to disturb the electrophysiology and contractility of atrial hiPSC-CMs via LATS2 phospho-regulatory effects upon YAP signaling. Supplementation of the SYNPO2L_A isoform can restore many functional deficits and supports a specific gene regulatory role for SYNPO2L in modulating the risk of AF and heart failure.

## INTRODUCTION

The gene SYNPO2L (synaptopodin like 2) is a striated muscle-specific protein localized to the Z-disk and implicated experimentally in the regulation of actin cytoskeleton organization and cardiac contractility^1,2^. Unlike its paralog SYNPO2, which is highly expressed in skeletal myocytes and non-striated muscle cells where it interacts with α-actinin and filamin C to mediate actin reorganization in myofibril assembly^3,4^, existing human single data suggest SYNPO2L is expressed primarily in cardiomyocytes. The gene structure of SYNPO2L is conserved to teleost fish^1^ and encodes two isoforms: a longer isoform SYNPO2L_A that is distinguished by the presence of an N-terminal PDZ domain, and a shorter isoform SYNPO2L_B that lacks the PDZ domain.

In a national biobank setting originating from Finland, a rare genetic variant predicted to interrupt splicing of the longer SYNPO2L_A isoform was observed to confer more than 3-fold increased risk for atrial fibrillation (AF)^5^. Additionally, common genetic variation in and around SYNPO2L has been strongly implicated in AF, electrocardiogram (ECG) phenotypes, and heart failure^6–9^. Epigenetic analyses of post-mortem human left atrial tissues from individuals without AF highlight a role of SYNPO2L within a network of genes governing atrial biology^10^, which is further supported by independent analyses of known methylation quantitative trait loci (mQTL) of SYNPO2L^11^. When combined with analyses of gene expression quantitative trait loci (eQTL) and splicing quantitative trait loci (sQTL) of SYNPO2L^5,12^, these findings together strongly implicate a fundamental role for the expression and splicing of SYNPO2L in atrial biology and risk of AF.

Here we describe deep characterization of a rare Finnish splicing mutant which specifically ablates the expression of SYNPO2L_A isoform to characterize the roles of SYNPO2L upon contractility and electrophysiological parameters in a human induced pluripotent stem cell derived cardiomyocyte (hiPSC-CM) model system. We observe changes to intracellular signaling suggesting specific mechanisms of SYNPO2L in regulating atrial cardiomyocyte biology along with downstream effects on gene expression. Phenotypic defects specific to atrial hiPSC-CMs are rescued by restoration of SYNPO2L_A via viral delivery, analogous to human cardiac gene therapies. Finally, analyses of large-scale human genetic data suggest a protective role for increased expression of SYNPO2L_A in a specific common form of post-ischemic AF.

## METHODS

### Generating Finnish Splice Mutant hiPSCs

WTC-11 hiPSCs with integrated blasticidin resistance at the MYH6 locus (WTC-BSD15) were used as the parental line for all hiPSCs in this study. CRISPR Cas9 mediated editing was used to introduce a homozygous knock-in SYNPO2L C>A mutation at chr10:73,655,817 (guide RNA sequence: CGUUACAGGUGUCUAAGGUA) of (gene name) in WTC-BSD15 hiPSCs by EditCo Bio, Inc. (Redwood City, CA, USA). To generate these cells, Ribonucleoproteins containing the Cas9 protein and synthetic chemically modified guide RNA produced by Synthego were electroporated into the cells using EditCo’s optimized protocol. Editing efficiency is assessed upon recovery, 48 hours post electroporation. Genomic DNA is extracted from a portion of the cells, PCR amplified and sequenced using Sanger sequencing. The resulting chromatograms are processed using EditCo’s Inference of CRISPR edits software (ice.synthego.com). To create monoclonal cell populations, edited cell pools are seeded at 1 cell/well using a single cell printer into 96 or 384 well plates. All wells are imaged every 3 days to ensure expansion from a single-cell clone. Clonal populations are screened and identified using the PCR-Sanger-ICE genotyping strategy described above. Only the clones containing the appropriate mutation frozen down and subsequently cultured for experiments, with mock-transfected WTC-BSD15 hiPSCs used as isogenic controls.

### hiPSC Culture and Maintenance

hiPSCs were cultured on Matrigel (Corning)-coated plates in mTeSR Plus (Stem Cell Technologies) at 37°C and 5% CO_2_ and medium was changed daily until cells achieved a confluency of approximately 80%. Cells were then passaged using 0.5 mM EDTA in PBS (Invitrogen) as needed until a sufficient quantity was obtained for differentiations.

### Differentiation of hiPSCs into Ventricular Cardiomyocytes

Cardiomyocytes were differentiated using a Wnt-modulating protocol^13^. In brief, when hiPSCs achieved a confluency of 70-80%, cells were induced with RPMI+B27(-insulin) (Invitrogen) + 7 μM CHIR990921 (LC Labs). Two days post-induction, cells were placed in RPMI+B27(-insulin) for 24 hrs, then in RPMI+B27(-insulin) supplemented with 5 μM IWP-2 (Sigma-Aldrich) for 48 hours. At 7 days post-induction, cells are then fed with RPMI+B27. At 9 days post-induction, cells are fed RPMI+B27 + 1 μM blasticidin (Gibco) to select for differentiated cardiomyocytes and these feeds last for 9 days with cells fed on alternative days. Purified cells were frozen down in CryoStor10 (StemCell Technologies). Only cardiomyocytes >90% cTnT^+^ as verified by flow cytometry were used in this study.

### Differentiation of hiPSCs into Atrial Cardiomyocytes

Differentiation of hiPSCs into atrial cardiomyocytes followed the same protocol as ventricular differentiation except for the addition of 1 μM retinoic acid (Sigma Aldrich) 3 days post-induction for 72 hrs as described^14^. Atrial identity of differentiated cardiomyocytes was verified by immunoblotting.

### qPCR

RNA was extracted from hiPSC-CMs using miRNAeasy MiniKit (Qiagen) and cDNA was synthesized using Superscript III First-Strand Synthesis (Invitrogen). Detection of mRNA was completed by using Taqman (Thermo Scientific) probes specific to human SYNPO2LA and SYNPO2LB. RT-qPCR was completed using a Quant Studio 7 Flex (Applied Biosystems). Human GAPDH served as a control for these studies.

### Immunoblotting

Protein was extracted from hiPSC-CMs using Pierce Ripa Buffer (Thermo Scientific) and 1:100 Halt Protease Inhibitor Cocktail (Thermo Scientific). Protein was quantified using a Pierce BCA Protein Assay Kit (Thermo Scientific) and loaded onto 4-12% Bis-Tris gels at 13 µg for SYNPO2L isoforms and at 5 µg for all other proteins. The higher loading mass for the SYNPO2L isoforms were due to lower expressional levels seen in hiPSC-CMs. Gels were run at 100 V for 1 hour before being transferred using an iBlot 3 Western Blot Transfer System (Thermo Scientific). The transferred blot was then blocked using Intercept Blocking Buffer (Licor). Blots were first incubated at 4°C overnight in primary antibody diluted in blocking buffer. The next day, the blot was washed using 1% TBS-T wash buffer and incubated for two hours in secondary antibody at room temperature. Blots were washed again in 1% TBS-T before being imaged using an Odyssey Imager (Licor). Specific antibodies used and their respective dilutions can be found in Table S1. Polyclonal antibodies against SYNPO2L_A and SYNPO2L_B isoforms were produced in rabbits by Genscript. For the A isoform, a custom protein consisting of amino acids 1-256 from the original SYNPO2L_A sequence was used to immunize the rabbits. For SYNPO2L_B isoform, rabbits were immunized with a small peptide containing 14 amino acids with homology to the full protein. The polyclonal antibodies produced were later purified and validated with protein lysates from hiPSC-CMs.

### Immunofluorescent Staining and Imaging

Atrial WTC cells were seeded in RPMI+B27+ 10% fetal bovine serum (FBS; Gibco) +1% Penicillin/Streptomycin (P/S; Gibco) +1 μM ROCKi (Biorbyt) on 96-well flat-bottom cell culture microplates (Greiner). After 24 hours, media was changed to fresh RPMI+B27+1% P/S and cells were maintained in this media for 7 days with media changes taking place every 2-3 days. 7 days post-seeding, cells were fixed using 4% paraformaldehyde for 15 minutes and blocked using 4% BSA+PBS-T solution for 1 hour. Primary antibodies were diluted at a 1:200 dilution in BSA+PBS-T and added to fixed cells for an overnight incubation. Following overnight incubation, cells were washed with PBS and secondary antibodies were diluted in 4% BSA+PBS-T at a 1:500 dilution and added onto cells for 2 hours. After 2 hours, cells were washed with PBS and imaged with a 63x oil immersion objective lens using a confocal microscope (Leica Stellaris 5).

### Immunoprecipitation

HEK293 T Cells were cultured in DMEM media supplemented with 10% fetal bovine serum (FBS). Plasmid transfection was performed using ViaFect (Promega) in Opti-MEM (Gibco) media according to the manufacturer’s instructions. 72 hours after transfection, cells were collected and then lysed with immunoprecipitation (IP) buffer (25 mM Tris, 150 mM NaCl, 1 mM EDTA, 1% NP40, 5% glycerol) with a cocktail of protease and phosphatase inhibitors. For FLAG magnetic bead immunoprecipitation, equal amounts of protein were incubated with magnetic beads that were initially bound to FLAG (MA1-91878, Thermo Scientific) or control (MA514447, Thermo Scientific) antibodies overnight at 4°C according to the Pierce Crosslink Magnetic Kit protocol (Thermo Scientific). Samples were washed three times with IP buffer before being resolved by sodium dodecyl sulphate-polyacrylamide gel electrophoresis (SDS-PAGE) and immunoblotted.

### AAV Production

AAV production was carried out as previously described^15^. Briefly, HEK293T cells were seeded in a HyperFlask Corning) and triple-transfected using a 2:1 polyethylenimine (PEI):DNA ratio with: a helper plasmid containing adenoviral elements (pHelper), a plasmid containing Rep2 and Cap genes from a proprietary AAV9 variant, CR9-01, that has a higher iPSC-CM transduction efficiency than parental AAV9 (U.S. Patent No.: US 2023/0220014 A1), and a third plasmid containing the SYNPO2L_A or No ORF control cassette to be packaged. 72 hours following transfection, cells were harvested and lysed. The virus was purified using iodixanol ultracentrifugation and was cleaned and concentrated in Hank’s Balanced Salt Solution (HBSS) with 0.001% Pluronic using a 100-kDa centrifuge column (Amicon). Purified AAV was titered using a PicoGreen assay and stored at −80°C until use.

### Electrophysiology Assessment with Multi Electrode Arrays

The MaestroPro (Axion Biosystems) microelectrode array instrument was used to collect electrophysiological measurements. Atrial WTC-Mock Transfected and atrial *SYNPO2L*^SS/SS^ cells were seeded in RPMI+B27+ 10% fetal bovine serum (FBS; Gibco) +1% Penicillin/Streptomycin (P/S; Gibco) +1 μM ROCKi (Biorbyt) on 24-well Cytoview plates coated with 98 μg/mL Matrigel (Corning). After 24 hours, media was changed to fresh RPMI+B27+1% P/S and cells were maintained in this media for 14 days with media changes taking place every 2-3 days. Cells were then placed in cardiac maturation media^16^ on day 14. On day 15 and day 18 post-seeding, cells were paced at 2.5 Hz for 1 hour and for 0.5 hours, respectively, to induce further maturation. Baseline electrophysiological readouts were taken at day 28. Two days later, a 5K multiplicity of infection (MOI) dose of AAV:SYNPO2L_A was added to the cells for 24 hours. Readouts were taken at 20 days post-AAV transduction. The first 30 seconds of a 3-minute measurement was used to analyze both field potential and action potential measurements. All data was batch processed using the AxIS Navigator 3.1.12 (Axion BioSystems) and analyzed with the Cardiac Analysis Tool 3.3.1 (Axion BioSystems). For field potential measurements, the T-wave peak detection was selected and verified manually by a blinded operator using methods as described by the manufacturer. For action potential analysis, two blinded operators selected a trace which was clearly captured and demonstrated a defined depolarization, peak, repolarization and return to resting membrane potential. Final computation of all readouts was performed by the Cardiac Analysis Tool 3.3.1. The tool corrected APD_90_ for beat rate according to Fridericia’s method and provided corrected measurements as APDc_90_.

### Live Cell Calcium Imaging

Calcium transients were recorded and analyzed utilizing the Nautilai medium-throughput calcium imaging instrument (CuriBio). 3×10^5^ cells were seeded in RPMI+B27 + 10% FBS + 1% P/S (Gibco) on Matrigel-coated 24-well plates (Corning). Cells were maintained in RPMI+B27 for 21 days before imaging. A working solution of Cardiomyocyte Maintenance Media (Cellular Dynamics) containing 5 μM of Cal520 dye (AAT Bioquest) and 0.04% Pluronic F-127 (Invitrogen) was added to cells and incubated at 37°C for 1 hour. Media was then aspirated and replaced with fresh maintenance media immediately prior to recordings Data was analyzed using the Pulse Cloud Platform (Curi Bio).

### Transcriptional Analysis by RNA Sequencing

Extraction of RNA was conducted using the same protocol as for qPCR with an additional step of DNA digestion using the RNase-Free DNase Set (Qiagen). The purified RNA was then sent out to Seqmatic for library preparation and RNA sequencing. From each replicate, 100 ng total RNA was extracted via the llumina Stranded mRNA Library Prep. RNA quality control was performed before library preparation using Agilent TapeStation instrument. The RNA libraries were prepared using a Stranded Total RNA Library Prep with Ribo-Zero Plus kit (Illumina), which also removes ribosomal RNA. The libraries were sequenced as 2×150 base pair paired-end reads using Illumina NovaSeq X Plus using a 10B flow cell with an average of 25 million reads per each read file. After adapter trimming by *fastp* (version 0.23.3), raw RNA-seq reads from hiPSCs in fastq format were aligned with *Salmon* (version 1.8.0) to the GENCODE (version 38, May 2021) reference transcript assembly (GRCh38.p13 and Ensembl 111) using best practice parameters to ensure mapping validity and reproducibility (--seqBias --gcBias --posBias --useVBOpt --rangeFactorizationBins 4 --validateMappings --mimicStrictBT2). Next, a script using R package *tximport* was used to generate an expression matrix normalized to transcripts per million (TPM) per gene and per isoform transcript. In this analysis, we only used genes detected in at least 10% of all samples. Protein-coding genes were determined using Ensembl Homo sapiens annotations (release 111, Jan 2024) and extracted by *biomaRt* (version 2.52.0). Mitochondrial genes were also excluded, followed by renormalization to TPM. These gene expression values were then log2-transformed after addition of 1 as pseudo-count. Expression patterns of key genes associated with functions of interest were visualized across treatment groups with boxplots generated using the *ggplot2* R package. Relative gene expression levels across groups are also presented in scaled values per gene in the heatmaps. Heatmaps were generated in R using *ComplexHeatmap* package.

For initial assessment and identifying presence of cluster patterns in the transcriptome, Principal Component Analysis (PCA) models were generated in R using the ‘*prcomp*’ function from the stats package. The first two principal components were used to visualize group level differences across samples in a PCA plot generated using ggplot2 and ggfortify packages with the ‘autoplot’ function. Differential gene expression analysis was then performed by comparing each two groups of interest using Welch’s *t*-test on pseudo-log normalized TPM values. The obtained t statistics values were used to rank-order the genes for the downstream functional analyses. Volcano plots were then generated to visualize the top positive and negative differentially expressed genes (DEGs) using the ggplot2 R package. The top DEGs are the set of genes with the highest and lowest t-statistics values. To evaluate functional effects, we performed Gene Set Enrichment Analysis (GSEA)^17^ on the gene list pre-ranked by t-statistics obtained from differential gene expression analysis, using the *clusterProfiler* R package. GSEA assesses whether differences in expression of predefined gene sets between two phenotypes are concordant and statistically significant. Gene sets were obtained from positional, curated canonical pathways, transcription factor targets, gene ontology, cell type signatures and Hallmark collections in Human MSigDB (v2023.1.Hs)^18^. Upon performing GSEA, these gene sets were only considered statistically significant if the false discovery rate (Q value) was less than 0.25 as determined with multiple hypothesis testing correction using the BH-correction method. The normalized enrichment score, which reflects the degree to which a gene set is overrepresented in the ranked list and normalized for gene set size, was used to select significantly altered gene sets.

### Engineered Heart Tissues

Engineered heart tissues (EHTs) were fabricated using the commercially available Mantarray kit (Curi Bio), and manufacturer’s protocols were used. In brief, casting wells arrayed in a 24-well plate format were pre-filled with 50 μL of a 6.4 U/mL thrombin solution, and the post lattice was inserted into these wells. hiPSC-CMs and human cardiac fibroblasts (HCFs) were then mixed with 5 mg/mL of fibrinogen at a composition of 5×10^5^ cardiomyocytes and 7.5×10^4^ HCFs per tissue. This cell and fibrinogen solution was then thoroughly mixed with the thrombin around the inserted posts. Mixtures were allowed to crosslink for 80-90 minutes at 37°C before being lifted out of the casting wells and the now-formed tissues placed into a new 24-well plate filled with fresh RPMI+B27 media. EHTs typically began beating within one week of casting. At 7 days post-casting, EHTs were cultured in maturation media^16^ for the duration of the experiment. Transduction with 5K MOI AAV:No ORF and AAV:SYNPO2L_A occurred at 11 days post-casting. Endpoint measurements were taken 20 days post-transduction with the EHTs paced at 1.5 Hz. Collected raw data was processed via the manufacturer’s cloud-based software (Curi Bio Pulse3D) to produce contractility metrics.

### *In silico* UK Biobank Analyses of Atrial Fibrillation

The UK Biobank (UKB) is a prospective cohort study with genotype and detailed phenotype information for up to 502,628 individuals aged 40-69 years when recruited between 2006 and 2010. UKB participant data is enriched with demographic characteristics, lifestyle details, and self-reported health outcomes.

Propensity score matching, combined with thorough survival analysis, was meticulously conducted to elucidate the impact of the SYNPO2L rs60632610 variant on the temporal progression to atrial fibrillation diagnosis or severe cardiovascular outcomes subsequent to the initial diagnosis of ischemic heart disease. Participants underwent meticulous matching based on a comprehensive array of covariates, including sex, smoking status, C-reactive protein levels (measured at the time of initial assessment), body mass index (BMI), low-density lipoprotein levels (LDL) at the time of initial assessment, first measurement of systolic blood pressure, age at initial ischemic heart disease diagnosis, presence of type 1 or type 2 diabetes, chronic renal failure diagnosis, and medication utilization encompassing beta-blockers, angiotensin-converting enzyme (ACE) inhibitors, calcium channel blockers, vasodilator antihypertensive drugs, centrally acting hypertensives, alpha-adrenoceptor blocking drugs, statins, digoxin, angiotensin II receptor blockers (ARBs), diabetic medications, and ezetimibe. This meticulous approach ensures robust control of confounding factors, thereby enabling a nuanced examination of the association between the SYNPO2L rs60632610 variant and cardiovascular health outcomes in individuals with ischemic heart disease. More detailed information on matching and survival analysis procedures can be found in a recent preprint^19^.

Co-localization analysis was performed with Hypothesis Prioritisation in multi-trait Colocalization (HyPrColoc) which is an efficient deterministic Bayesian divisive clustering algorithm using GWAS summary statistics to detect colocalization across vast numbers of traits simultaneously^20^. In our analysis we tested left and right heart structure and function^21^, DCM^22^, atrial fibrillation^7^, systolic and diastolic blood pressure^23^ and heart failure from the Global Biobank Meta-analysis Initiative (GBMI)^24^. We specified a region of SYNPO2L ± 2Mb to compute the posterior probability that the traits share a causal variant.

### Statistical Analyses

Statistical analyses were performed with either the R language for statistical computing or with GraphPad Prism. A Student’s t-test was performed for all comparisons between only two groups, while one-way ANOVA with a Tukey’s post-hoc test was performed for all datasets with multiple group comparisons. To characterize the number of extra-systolic beats in paced data, a linear fit model accounting for time was used. Cox proportional hazards regression was used for survival curve analyses.

## RESULTS

### Finnish Splice Mutation Results in SYNPO2L_A Haploinsufficiency

The rare splice mutation is located at the 3’ splice junction of the 1^st^ exon and is predicted to disturb splicing of the PDZ domain encoded by the first two exons to encode the SYNPO2L_A isoform (Fig. 1A). To examine the effect of this mutation on cardiomyocytes (CMs) *in vitro*, we created an hiPSC line which was homozygous for the rare Finnish splice mutation, referred to hereafter as the splice mutant (*SYNPO2L*^SS/SS^), and differentiated this line and a isogenic wild-type control (WTC) line into high-purity ventricular or atrial hiPSC-CMs using established protocols^13,14^. Differentiated atrial CMs displayed standard markers of atrial phenotypes^14,25^ including decreased expression of MLC2v, significantly shorter action potential durations (APDs), reduced contractility, and increased spontaneous beat rates (Fig. S1). Western blotting of WTC cardiomyocytes shows that a significantly higher ratio of SYNPO2L_B to SYNPO2L_A protein was present in atrial hiPSC-CMs compared to ventricular hiPSC-CMs (Fig. S2). Expression of *SYNPO2L_A* RNA was found to be negligible in *SYNPO2L*^SS/SS^ ventricular cardiomyocytes compared to WTC, while *SYNPO2L_B* expression was slightly greater in mutant cells, and the same trend was observed with regards to SYNPO2L_A and SYNPO2L_B protein expression (Fig. 1C). Similarly, the introduction of the splice mutation in atrial CMs led to a significant decrease in SYNPO2L_A expression at both the transcript and protein level relative to WTC (Fig. 1D).

**Figure 1.**
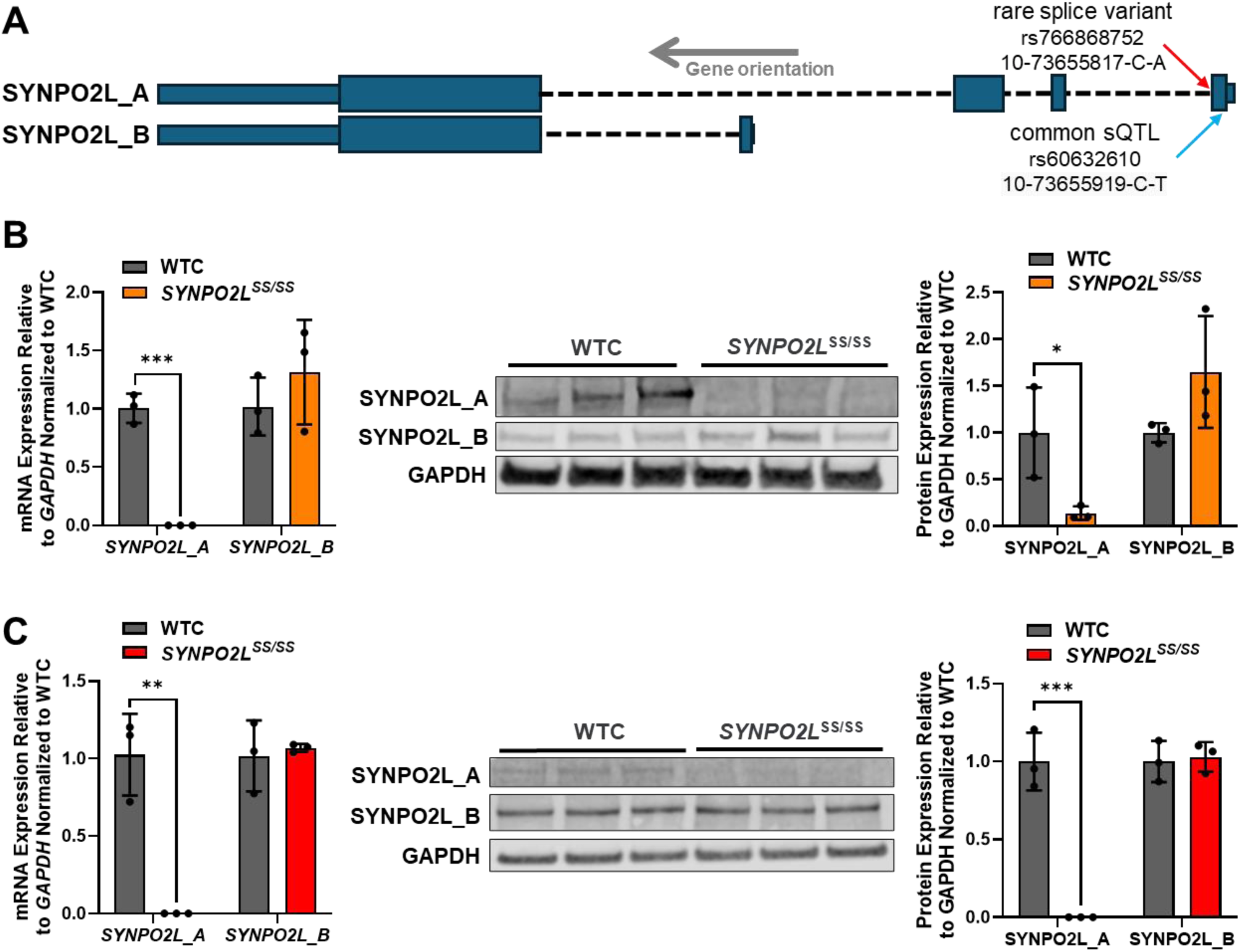
SYNPO2L splice mutation induces SYNPO2L_A haploinsufficiency. (A) Schematic detailing the location of the rs766868752 splice variant mutation on the genome and its relation to the two isoforms of SYNPO2L. (B) Analysis with qRT-PCR and western blot shows that expression of the SYNPO2L_A isoform is negligible at the transcript and protein levels in ventricular *SYNPO2L*^SS/SS^ hiPSC-CMs. A slight increase in SYNPO2L_B expression is also observed. (C) SYNPO2L_A expression is also negligible in atrial *SYNPO2L*^SS/SS^ hiPSC-CMs at the transcript and protein levels. n=3 biological replicates. *p<0.05, **p<0.01, ***p<0.001 as determined by Student’s t-test.

### Electrophysiological Perturbations are Induced by Splice Mutation

Given the linkage of SYNPO2L to atrial physiology and disease^26–28^, we examined the impact of the splice mutant upon electrophysiological properties using multi-electrode arrays (MEAs). By day 14 of culture, splice mutant atrial CMs exhibited distinct differences in their action potentials compared to WTC atrial CMs such as an extended plateau phase and a longer time to repolarization (Fig. 2A). Quantitative analysis confirmed significantly prolonged APDs in addition to lower field potential spike amplitudes and decreased spike slopes in comparison to WTC CMs (Fig. 2B). Increased APDs are indicative of QT prolongation, and the latter phenomenon has been closely linked to an increased risk of arrhythmias and atrial fibrillation^29^. To confirm this observation, we performed an extended pacing of the cells at 2.1 Hz for a total of 90 minutes with 3 minute long recording periods every 15 minutes, and we observed that not only did *SYNPO2L*^SS/SS^ atrial cardiomyocytes exhibit greater beat period variation (Fig. S3A), but that the mutant atrial hiPSC-CMs displayed aberrant beats in 23.8% of timepoints measured, compared to no aberrant beats produced by the WTC control atrial hiPSC-CMs (Fig. S3B, p=0.007). Notably, an increased frequency of premature atrial contractions is a recognized risk factor for atrial fibrillation^30^. We additionally measured intracellular calcium dynamics in WTC and *SYNPO2L*^SS/SS^ atrial cardiomyocytes during spontaneous beating using live imaging of Fluo-4 AM, a fluorescent Ca^2+^-sensitive dye, and observed not only decreased rates of calcium influx and reuptake, but an overall decrease in calcium transient amplitude as well (Fig. 2C). For comparison, splice mutant ventricular CMs displayed only mild differences in the same electrophysiological and calcium handling parameters (Fig. S4).

**Figure 2.**
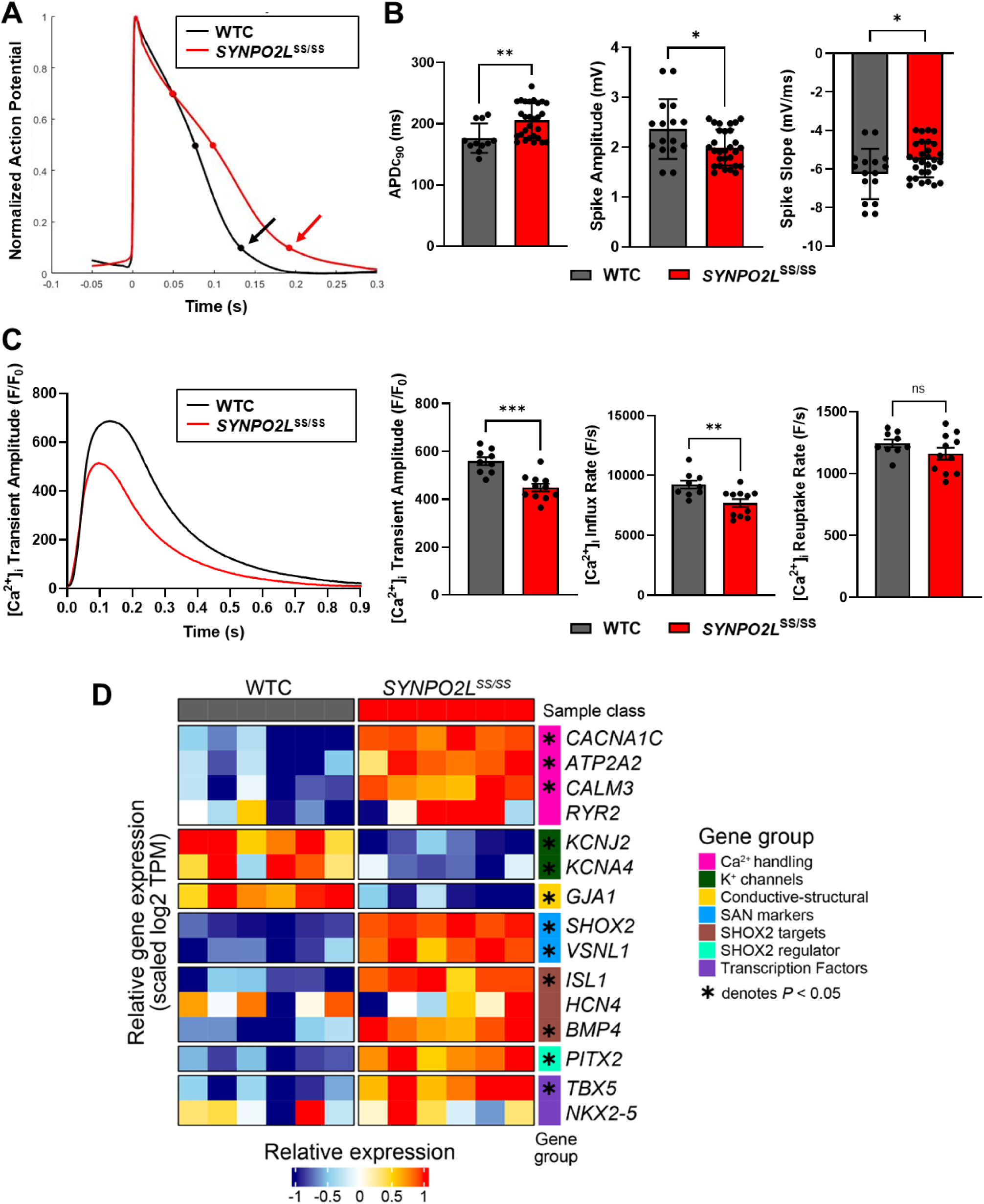
Electrophysiological function is perturbed in *SYNPO2L*^SS/SS^ atrial hiPSC-CMs. (A) Representative action potential traces illustrating the prolonged APD_90_ in *SYNPO2L*^SS/SS^ cells (red arrow) compared to WTC cells (black arrow). (B) Significant increases in APDc_90_ in mutant atrial hiPSC-CMs is observed, as well as significant decreases in both field potential spike amplitudes and spike slopes. (C) Representative trace of calcium transients in WTC and *SYNPO2L*^SS/SS^ cells. Quantification of fluorescent calcium imaging shows significant decreases in transient amplitudes and influx rates, and slight decreases in reuptake rates in mutant cardiomyocytes. (D) RNA-seq analysis shows decreased expression of key ion channel and conduction genes, while expression of calcium handling, sinoatrial node (SAN), and *SHOX2* related genes are increased. n≥6 biological replicates in MEA data; n≥9 biological replicates in calcium imaging data n=6 biological replicates in RNA-seq data. *p<0.05, **p<0.01, ***p<0.001 as determined by Student’s t-test for MEA and calcium data; *p<0.05 as determined by Welch’s t-test for RNA-seq data.

We additionally performed RNA sequencing to characterize the transcriptional response of the splice mutation in *SYNPO2L*^SS/SS^ atrial CMs (Fig. 2D). A notable downregulation was observed of *KCNJ2* and *KCNA4*, both of which code for potassium ion channels that are key regulators of membrane potential repolarization and are known to increase arrhythmic risk when their function is disrupted^31,32^. Additionally, several critical Ca^2+^ handling genes were found to be significantly upregulated, including *CACNA1C*, *ATP2A2*, and *RYR2*. Similar to the aforementioned potassium channels, these calcium channels and receptor play important roles in maintaining proper excitation-contraction coupling and aberrations in their expression or function have been demonstrated to lead to arrhythmogenic events^33,34^. *GJA1* was also found to be downregulated in *SYNPO2L*^SS/SS^ cells. Dysregulation or remodeling of Cx43 has been observed in patients with AF^35^ and in animal models of AF^36^, and overexpression of Cx43 in large animal studies has been found to maintain atrial conduction and prevent persistent AF^36^. Interestingly, the most differentially upregulated gene between wild-type and mutant cells was *SHOX2*, a notable sinoatrial node (SAN) marker^37,38^. Correspondingly, multiple *SHOX2* targets (*ISL1*, *BMP4*)^39^, regulators (*PITX2*)^40^, and interacting transcription factors (*TBX5*) were also upregulated in mutant cells, as well as *VSNL1*, another SAN marker^41,42^. This abnormal shift in cardiomyocyte identity towards one of a pacemaker-like cell may therefore have also contributed to the electrophysiological dysfunction strongly consistent with a pro-arrhythmic phenotype that was observed in the *SYNPO2L*^SS/SS^ atrial cardiomyocytes.

### Impaired Contractility and Increased Arrhythmogenicity in *SYNPO2L*^SS/SS^ Tissues

To assess potential differences in contractile properties, WTC and *SYNPO2L*^SS/SS^ atrial hiPSC-CMs were formed into engineered heart tissues (EHTs) and cultured for 21 days prior to assessment. During both spontaneous beating and when paced at 1.5 Hz, splice mutant atrial hiPSC-CMs displayed dramatically lower twitch force and contraction velocities, and notably, greater beat-to-beat variations in twitch force and timing, which are both indicative of arrhythmic behavior (Fig. 3A)^43,44^. Important differences were also observed in the diastolic properties of splice mutant atrial hiPSC-CM EHTs, where *SYNPO2L*^SS/SS^ EHTs exhibited significantly decreased relaxation velocities and values for RT_90_ (Fig. 3B). In contrast, EHTs comprised of ventricular hiPSC-CMs carrying the splice mutant displayed more subtle variations in contractile function compared to WTC under both un-paced and paced conditions (Fig. S5). Transcriptomic analysis of the atrial EHTs showed dramatic downregulation of *TTN* and *ACTN2* in the mutant tissues (Fig. 3C). In combination with the electrophysiological readouts, the EHT data overall suggests that functional deficits may arise from the haploinsufficiency of a specific SYNPO2L isoform, and that the severity of these effects may furthermore be heart chamber specific.

**Figure 3.**
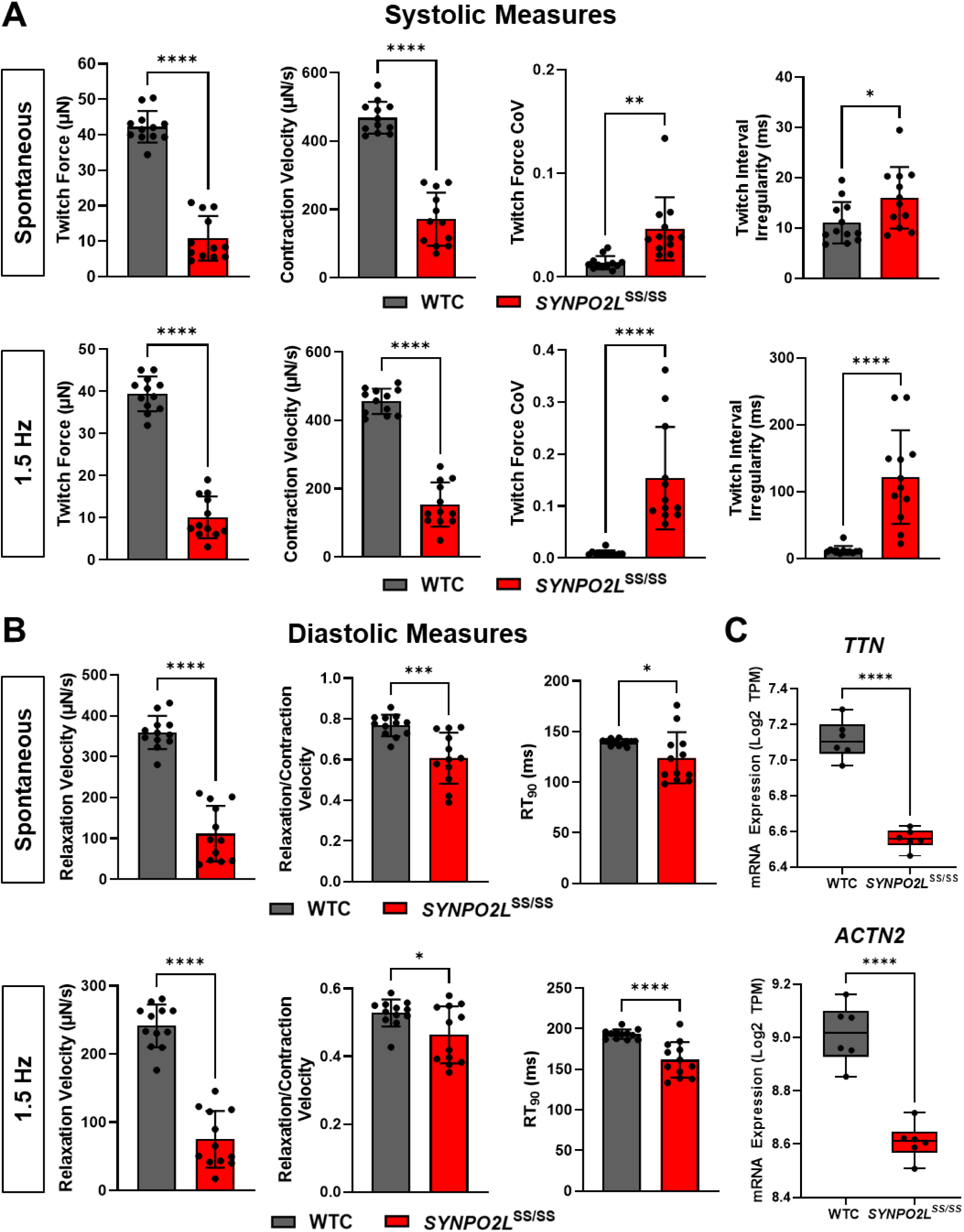
Impaired contractility of *SYNPO2L*^SS/SS^ atrial EHTs. (A) Under both spontaneous beating and paced conditions, *SYNPO2L*^SS/SS^ EHTs display decreased twitch forces and contraction velocities while measures for arrhythmic contraction such as twitch force coefficient of variance (CoV) and twitch interval irregularity are significantly increased. (B) Diastolic measures of contractility were also impacted, with mutant EHTs exhibiting reduced relaxation velocities, relaxation/contraction velocity ratios, and relaxation times. (C) Transcriptomic analysis revealed decreased expression of *TTN* and *ACTN2*, two important genes for contractile function regulation, in mutant EHTs. n=12 biological replicates. *p<0.05, **p<0.01, ****p<0.0001 as determined by Student’s t-test.

### SYNPO2L_A Modulates YAP Signaling via LATS2 Stabilization

The paralog of SYNPO2L has been observed to interact with the LATS1/2 complex to modulate YAP/TAZ signaling in other model systems^45^, and the role of YAP/TAZ in the HIPPO signaling pathway in cardiac development and disease is well recognized^46–48^. Therefore, we hypothesized that SYNPO2L_A might also interact with the LATS1/2 complex to modulate YAP/TAZ signaling in hiPSC-CMs (Fig. 4A)^49,50^. Co-localization experiments confirmed that SYNPO2L was localized to the α-actinin-rich Z-disks and that punctate LATS2 staining was present along the sarcomere in between the Z-disks (Fig. 4B), suggesting organelle-specific spatial restriction and interaction within the sarcomere, but not direct evidence of co-localization within a protein complex between LATS2 and SYNPO2L_A. To further interrogate this observation, co-immunoprecipitation pull-down with an anti-FLAG antibody was performed with lysates from HEK cells that expressed LATS2 and were transfected with FLAG-tagged SYNPO2L_A. A significantly stronger band for LATS2 was observed in the eluted FLAG-tagged sample compared to the input, indicating that LATS2 was bound to SYNPO2L_A and that an interaction occurs between the two proteins in mammalian cells (Fig. 4C). Interestingly, while assessment of *LATS1* and *LATS2* transcript expression in atrial cardiomyocytes suggested no appreciable difference between WTC and *SYNPO2L*^SS/SS^ cells (Fig. 4D), immunoblotting showed that at the protein level, LATS2 expression was significantly decreased in the splice mutant cells (Fig. 4E), supporting the hypothesis that the lack of SYNPO2L_A diminishes the stability of LATS2. Immunoblotting also suggested that while YAP levels were indistinguishable between the splice mutant and WTC cardiomyocytes, phosphorylated YAP (pYAP) was significantly lower in the splice mutant, and this decrease in phosphorylation is consistent with the observed instability of LATS2 (Fig. 4F). Additionally, in transcriptional analyses of WTC and *SYNPO2L*^SS/SS^ EHTs, we noted broad and significant changes in known downstream targets of YAP transcriptional modulation (Fig. 5, Table S2). Together, these data may support a SYNPO2L isoform specific effect upon binding LATS2 to modulate downstream targets of YAP/TAZ.

**Figure 4.**
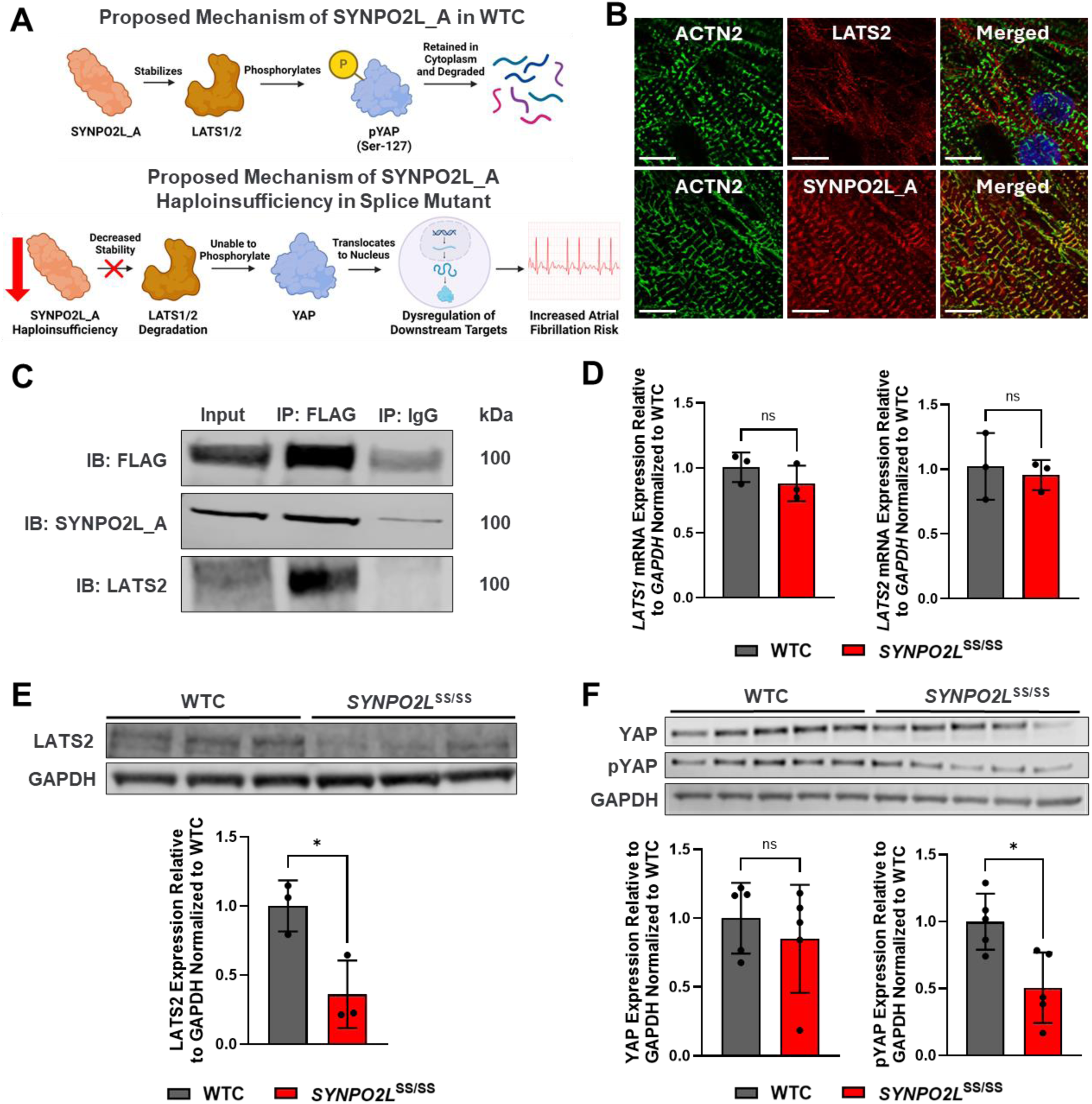
SYNPO2L_A acts to stabilize LATS2, a regulator of YAP phosphorylation. (A) Schematic detailing the hypothesized function of SYNPO2L_A as a stabilizer of LATS1 and LATS2 which allows for the phosphorylation of YAP to occur, and how this mechanism is impacted by the haploinsufficiency of SYNPO2L_A in splice mutant cells. (B) Representative immunofluorescence images of atrial hiPSC-CMs stained for ACTN2, LATS2, and SYNPO2L_A. SYNPO2L_A was found to be co-localized to sarcomeres. Scale bar = 10 μm. (C) Co-immunoprecipitation of FLAG-tagged SYNPO2L_A and LATS2 produced a higher intensity band for LATS2, indicating that the two proteins bind and interact together in mammalian cells. (D) The splice mutation does not impact the transcriptional expression of both *LATS1* and *LATS2* as determined by qRT-PCR. (E) However, the western blots indicate that the quantity of LATS2 at the protein level was found to be significantly decreased in *SYNPO2L*^SS/SS^ atrial cardiomyocytes. (F) Although protein levels of YAP are comparable between WTC and mutant cardiomyocytes, the amount of phosphorylated YAP (pYAP) is significantly reduced in mutant cells. n≥3 biological replicates. *p<0.05 as determined by Student’s t-test.

**Figure 5.**
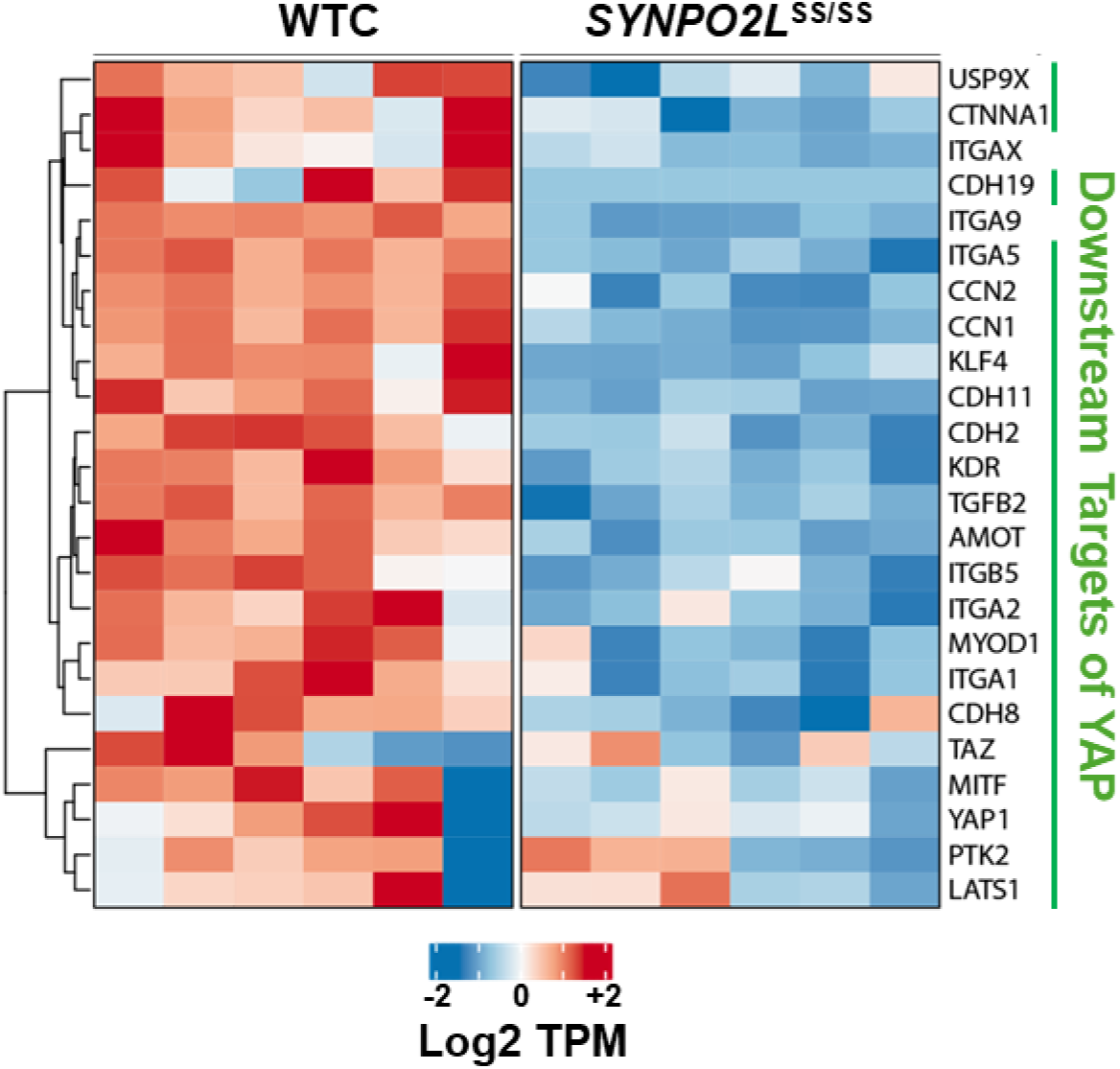
RNA sequencing illustrates transcriptional impact of perturbed YAP phosphorylation in mutant cardiomyocytes. A variety of downstream targets of YAP are downregulated in *SYNPO2L*^SS/SS^ atrial cardiomyocytes, suggesting that a reduction of YAP phosphorylation due to a loss of SYNPO2L_A leads to cascading effects at the transcriptional level. n=6 biological replicates.

### Diminished Pro-arrhythmic Dysfunction with SYNPO2L_A Overexpression

Next, we sought to examine the effects of restoring intracellular expression of SYNPO2L_A upon the pro-arrhythmic phenotype, abnormal calcium flux, and decreased contractility associated with reduced SYNPO2L_A levels in the splice mutant cell line. We constructed a vector with an expression cassette containing the SYNPO2L_A isoform under the control of the human TNNT2 promoter (AAV:SYNPO2L_A), along with a vector containing a cassette without an open reading frame (AAV:No ORF) (Fig. S6), and packaged each into a AAV9-derivative capsid. *SYNPO2L*^SS/SS^ hiPSC-CM monolayers were treated with virus after 30 days of culture with a 5K multiplicity of infection (MOI) dose of AAV:SYNPO2L_A for 24 hours. By 7 days post-transduction, expression of SYNPO2L_A in treated mutant cells had reached approximately 9-fold that of WTC (Fig. S7). MEA measurements taken 20 days post-transduction showed significant restoration of action potential durations, but only partial restoration of spike amplitudes and spike slopes, to levels observed in WTC cells treated with AAV:No ORF (Fig. 6A). Furthermore, the percentage of wells exhibiting extra beats under extended pacing was drastically reduced with AAV:SYNPO2L_A treatment, with a reduction of 21.4% observed (Fig. S8, p=0.006). In a parallel study with EHTs, treatment of splice mutant tissues with the same dose of AAV:SYNPO2L_A partially improved pre-treatment abnormalities in twitch force and contractile velocity, while more notably, metrics for arrhythmic contraction were fully normalized to WTC EHTs treated with AAV:No ORF (Fig. 6B). Diastolic function, in particular relaxation velocities and relaxation times, displayed some improvements towards WTC post-transduction, but the degree of rescue was not statistically significant at 20 days post-transfection (Fig. 6C).

**Figure 6.**
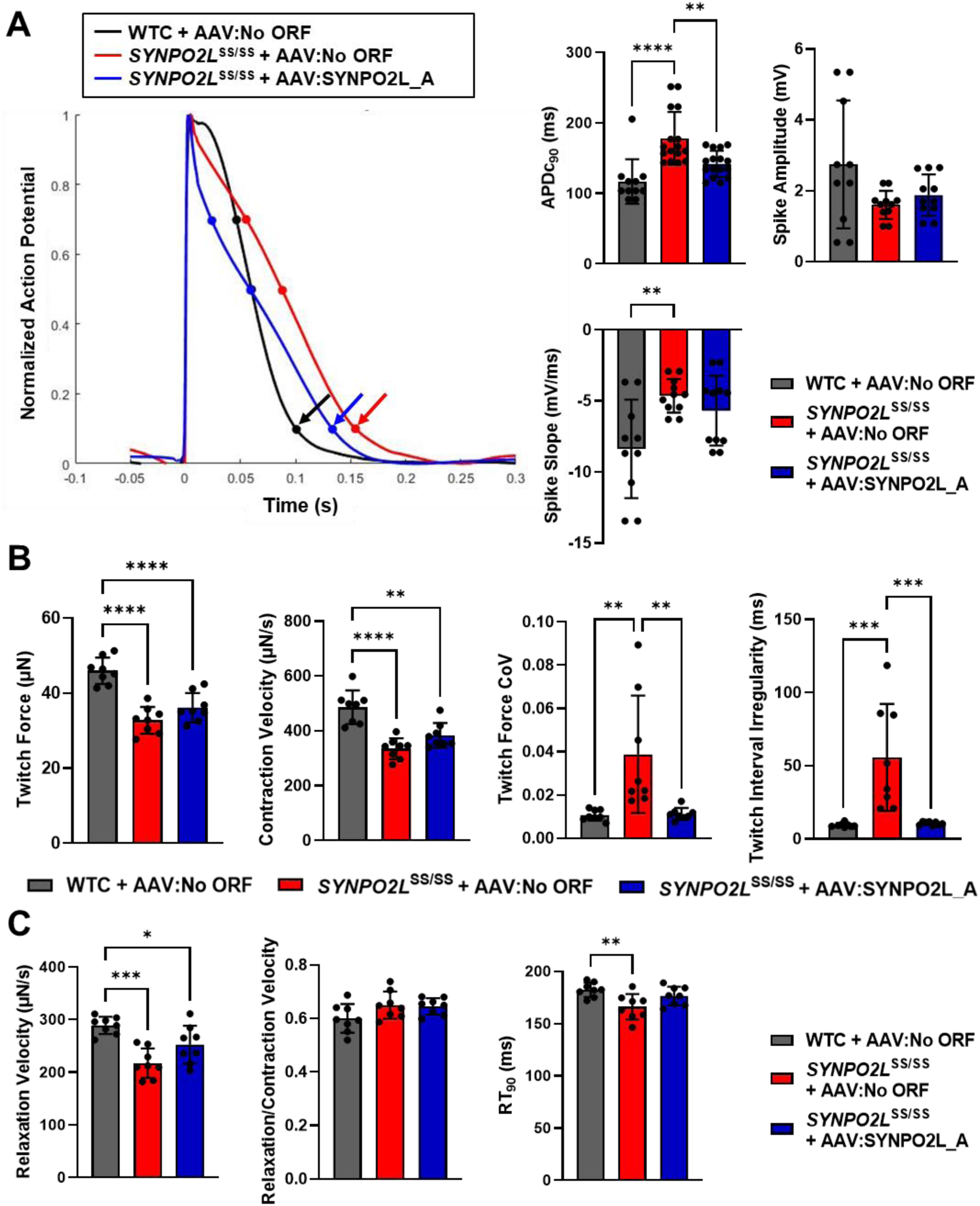
Overexpression of SYNPO2L_A results in normalization of splice mutation induced functional abnormalities in atrial hiPSC-CMs. (A) Action potential durations (APDs) are shifted leftwards with AAV treatment (colored arrows) and this is reflected in decreased APDc_90_ values in treated cardiomyocytes compared to AAV:No ORF controls. However, no significant changes in field potential spike amplitudes and spike slopes were observed post-treatment. (B) In EHTs, the addition of SYNPO2L_A resulted in slight increases in twitch forces and contraction velocities, but more drastic normalization occurred in metrics for arrhythmia such as twitch force CoVs and twitch interval irregularities. (C) Diastolic measures of contractile function in EHTs also showed improvements post-treatment in relaxation velocities and RT_90_ values, but no discernable changes were observed in the ratio between relaxation and contraction velocities. n≥6 biological replicates for MEA data; n=8 biological replicates for EHT data. *p<0.05, **p<0.01, ***p<0.001, ****p<0.0001 as determined by one-way ANOVA.

Given the observed efficacy of AAV:SYNPO2L_A in correcting the electrophysiological and contractile deficits observed in the splice mutant atrial hiPSC-CMs *in vitro*, we systematically examined groups of individuals affected with AF in the UK Biobank to develop a hypothesis about clinical scenarios which might also benefit from AAV:SYNPO2L_A supplementation. We performed an analysis using a splice quantitative trait locus (sQTL) for SYNPO2L rs60632610 where the T allele increases expression of SYNPO2L_A in the left atrium according to GTEx data^51^. Within the UK Biobank we defined three groups of individuals with a different AF origin, where a diagnosis of AF occurred more than 1 month after a primary diagnosis of: a) non-ischemic heart failure, b) mitral or aortic valve disease, or c) ischemic heart disease. Within each of these pre-defined groups, we used a recently described approach for matching individuals for age at diagnosis, sex, BMI, and medication use across the genotypes for the sQTL, thus establishing different ‘treatment’ groups who have higher or lower levels of SYNPO2L_A in the atrium. In this analysis we observed that individuals with the T allele, which increases the expression of SYNPO2L_A in the left atrium, appeared to be protective against AF specifically amongst individuals who experienced myocardial infarction (MI) (HR per allele 0.92, p=5.27e-6) (Fig. 7), but not in individuals experiencing AF after a diagnosis of valvular disease, nor in non-ischemic heart failure. Genetic co-localization analysis identified rs60632610 as a most likely causal SNP in the SYNPO2L region co-localizing with dilated cardiomyopathy, atrial fibrillation, heart failure, left ventricular ejection fraction, systolic and diastolic blood pressure and short axis pulmonary artery diameter (Table S3).

**Figure 7.**
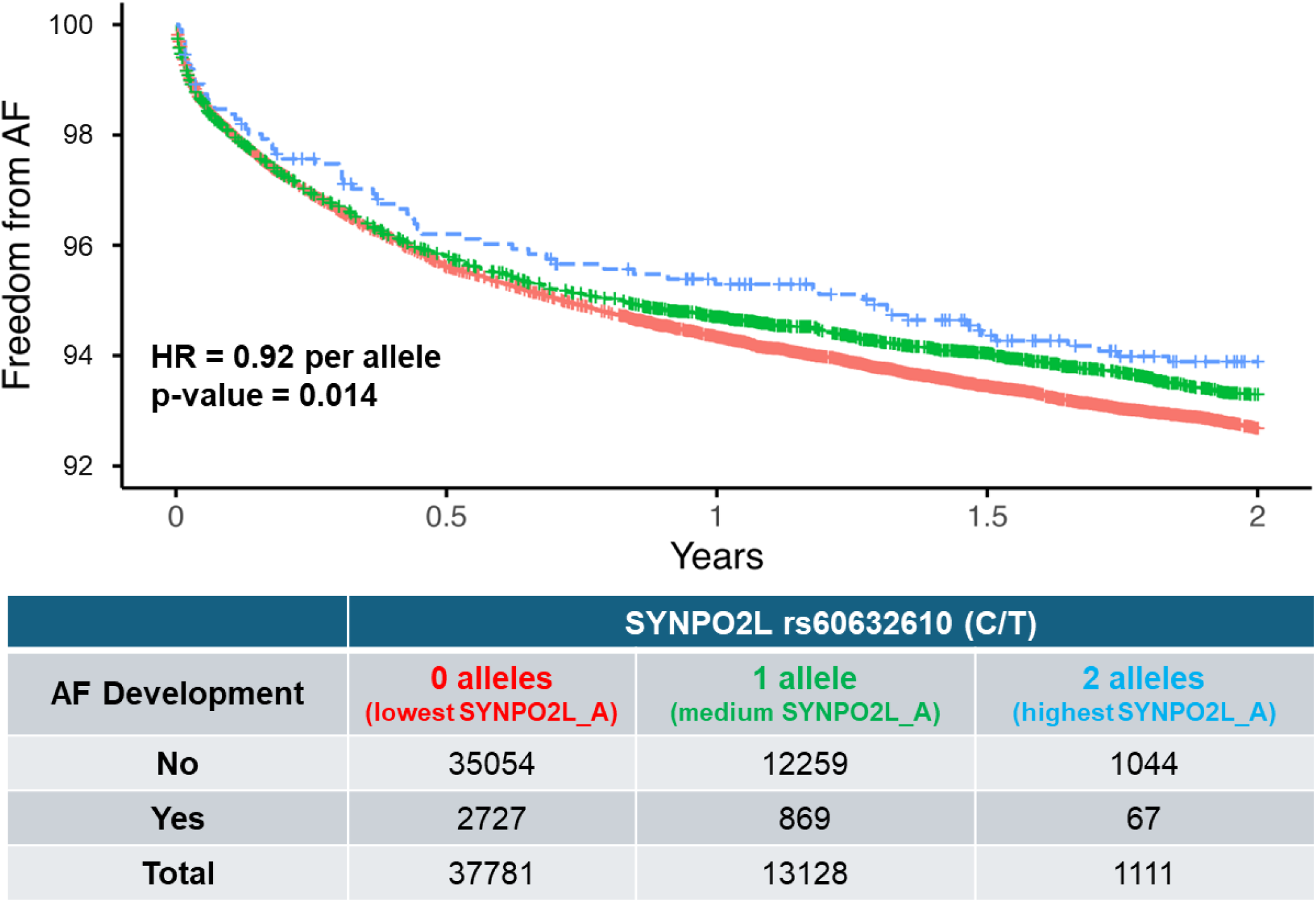
A common genetic variant increasing the level of SYNPO2L_A protects against development of atrial fibrillation after myocardial infarction. Effect of SYNPO2L variant rs60632610 on time to atrial fibrillation (AF) development from first ischemic heart disease diagnosis in UK Biobank. Carrying the T allele is associated with reduced risk of developing atrial fibrillation after ischemic heart diagnosis with HR of 0.92 and p=0.014 per allele as determined by Cox proportional hazards regression analysis.

## DISCUSSION

Here we demonstrate the functional consequences of human genetic variation in SYNPO2L where a rare splice mutant that ablates the SYNPO2L_A isoform confers risk for AF by compromising contractility and adversely impacting conduction and calcium handling specifically in atrial-hiPSC-CMs. Additionally, our findings suggest a molecular mechanism by which SYNPO2L plays an important role in atrial biology suggesting that the protein is localized to the Z-disk and the PDZ domain present in the A isoform but absent in the B isoform may interact with LATS2 to modulate YAP signaling. Our data suggests that exogenous restoration of SYNPO2L_A isoform can reverse *in vitro* the pathological electrophysiological and contractile abnormalities observed in deficient atrial hiPSC-CMs. Together these data elucidate a possible mechanism for SYNPO2L in atrial cardiomyocytes linking the sarcomere to modulation of pathogenic signaling modifying electrophysiological and contractile properties than change the risk of atrial arrhythmias.

Atrial fibrillation is an important and highly morbid complication affecting individuals after ischemic heart disease, and new approaches to treatment or prevention have the potential for impact upon cardiovascular health^52^. The use of human genetics to identify targets is widely accepted across the field of therapeutic discovery^53^, and the datasets and tools available have grown such that genetic approaches now have the potential to change risk stratification and patient selection in cardiovascular disease^54^. The value of studying rare genetic mutations, such as the splice mutant described in detail here, is that a potential patient population carrying this mutation (numbering likely in the hundreds) may also be simultaneously evaluated for a therapeutic intervention. As we describe in a related publication, careful use of instrumental variables derived from human genetics which are known to directly impact protein function, expression, or splicing, may permit the *in silico* evaluation of these proteins as therapeutic targets in specific clinical settings^19^.

In addition to atrial fibrillation risk, the role of SYNPO2L isoforms in cardiac development^2^, ventricular hypertrophy^10^ and heart failure^55^ are described and related to an actin binding where the A isoform facilitates nucleation of larger diameter actin bundles *in vitro*^56^. Besides a primary role stabilizing actin bundle formation, a gene regulatory role of SYNPO2L has also been suggested from *ex vivo* analyses of human atrial tissue^10^ which further supported by the LATS2 dependent mechanism suggested from the data presented here. Together these data suggest the possibility that SYNPO2L may impact cardiomyocyte function via both direct effects upon sarcomere structure and function alongside downstream transcriptional effects upon the YAP signaling.

Our study is not without limitations. The differentiation and culture conditions of atrial hiPSC-CMs *in vitro* likely cannot account for the decades of contractile cycles and shear stress, along with the millions of cardiac action potentials, which shape the *in vivo* maturation of atrial cardiomyocytes. In addition, the hiPSC-CM culture systems employed do not include immunological and endothelial cell types, endocrine stimulation, nor sympathetic and parasympathetic innervation, all of which contribute to normal physiology and pathophysiological disturbances relevant to multi-factorial diseases such as atrial fibrillation and heart failure. The splice site mutation was modeled in a homozygous cell line to elicit a clear phenotype, but individuals with increased risk of atrial fibrillation are heterozygous for this mutation. Some of the genetic techniques applied to survival analyses are untested for novel targets and indications in cardiovascular disease; therefore, the implications of these analyses may not be apparent until any potential therapies evaluated with this approach reach human populations.

Using atrial hiPSC-CMs, we have modeled the relationship of SYNPO2L splice isoforms to atrial contractility and electrophysiology and characterize a sarcomere-independent YAP/TAZ-dependent signaling mechanism by which the SYNPO2L_A isoform exerts a transcriptional effect on atrial biology. These findings demonstrate the utility of genetic models of hiPSC-CMs for understanding human biology and pathophysiology and may suggest modulation of SYNPO2L isoforms as a potential therapeutic approach to atrial fibrillation.

## ACKNOWLEDGEMENTS

The authors would like to thank Dr. Charles E. MacKay for his guidance with the co-immunoprecipitation experiments.

## SOURCES OF FUNDING

Funding for this study was provided entirely by Tenaya Therapeutics, Inc.

## DISCLOSURES

All authors were employees or contractors of Tenaya Therapeutics, Inc. at the time the work in this study was performed. All authors except for L.Z. and A.E.T. also hold equity in Tenaya Therapeutics, Inc.

## Notes

### Summary of Updates

Supplemental materials (figures and tables) have been added.

## REFERENCES

1. Beqqali A, Monshouwer-Kloots J, Monteiro R, Welling M, Bakkers J, Ehler E, Verkleij A, Mummery C, Passier R. CHAP is a newly identified Z-disc protein essential for heart and skeletal muscle function. J Cell Sci. 2010;123:1141–1150. doi: 10.1242/jcs.063859

2. van Eldik W, Beqqali A, Monshouwer-Kloots J, Mummery C, Passier R. Cytoskeletal heart-enriched actin-associated protein (CHAP) is expressed in striated and smooth muscle cells in chick and mouse during embryonic and adult stages. Int J Dev Biol. 2011;55:649–655. doi: 10.1387/ijdb.103207wv

3. Linnemann A, Vakeel P, Bezerra E, Orfanos Z, Djinović-Carugo K, Van Der Ven PFM, Kirfel G, Fürst DO. Myopodin is an F-actin bundling protein with multiple independent actin-binding regions. J Muscle Res Cell Motil. 2013;34:61–69. doi: 10.1007/s10974-012-9334-5

4. Linnemann A, van der Ven PFM, Vakeel P, Albinus B, Simonis D, Bendas G, Schenk JA, Micheel B, Kley RA, Fürst DO. The sarcomeric Z-disc component myopodin is a multiadapter protein that interacts with filamin and α-actinin. Eur J Cell Biol. 2010;89:681–692. doi: 10.1016/j.ejcb.2010.04.004

5. Clausen AG, Vad OB, Andersen JH, Olesen MS. Loss-of-function variants in the SYNPO2L gene are associated with atrial fibrillation. Front Cardiovasc Med. 2021;8:650667. doi: 10.3389/fcvm.2021.650667

6. Miyazawa K, Ito K, Ito M, Zou Z, Kubota M, Nomura S, Matsunaga H, Koyama S, Ieki H, Akiyama M, et al. Cross-ancestry genome-wide analysis of atrial fibrillation unveils disease biology and enables cardioembolic risk prediction. Nat Genet. 2023;55:187–197. doi: 10.1038/s41588-022-01284-9

7. Nielsen JB, Thorolfsdottir RB, Fritsche LG, Zhou W, Skov MW, Graham SE, Herron TJ, McCarthy S, Schmidt EM, Sveinbjornsson G, et al. Biobank-driven genomic discovery yields new insight into atrial fibrillation biology. Nat Genet. 2018;50:1234–1239. doi: 10.1038/s41588-018-0171-3

8. Levin MG, Tsao NL, Singhal P, Liu C, Vy HMT, Paranjpe I, Backman JD, Bellomo TR, Bone WP, Biddinger KJ, et al. Genome-wide association and multi-trait analyses characterize the common genetic architecture of heart failure. Nat Commun. 2022;13:6914. doi: 10.1038/s41467-022-34216-6

9. Verweij N, Benjamins JW, Morley MP, van de Vegte YJ, Teumer A, Trenkwalder T, Reinhard W, Cappola TP, van der Harst P. The genetic makeup of the electrocardiogram. Cell Syst. 2020;11:229–238. doi: 10.1016/j.cels.2020.08.005

10. Hall AW, Chaffin M, Roselli C, Lin H, Lubitz SA, Bianchi V, Geeven G, Bedi K, Margulies KB, De Laat W, et al. Epigenetic analyses of human left atrial tissue identifies gene networks underlying atrial fibrillation. Circ Genomic Precis Med. 2020;13:e003085. doi: 10.1161/CIRCGEN.120.003085

11. Liu Y, Li B, Ma Y, Huang Y, Ouyang F, Liu Q. Mendelian randomization integrating GWAS, eQTL, and mQTL data identified genes pleiotropically associated with atrial fibrillation. Front Cardiovasc Med. 2021;8:745757. doi: 10.3389/fcvm.2021.745757

12. Hsu J, Gore-Panter S, Tchou G, Castel L, Lovano B, Moravec CS, Pettersson GB, Roselli EE, Gillinov AM, McCurry KR, et al. Genetic control of left atrial gene expression yields insights into the genetic susceptibility for atrial fibrillation. Circ Genomic Precis Med. 2018;11:e002107. doi: 10.1161/CIRCGEN.118.002107

13. Lian X, Hsiao C, Wilson G, Zhu K, Hazeltine LB, Azarin SM, Raval KK, Zhang J, Kamp TJ, Palecek SP. Robust cardiomyocyte differentiation from human pluripotent stem cells via temporal modulation of canonical Wnt signaling. Proc Natl Acad Sci U S A. 2012;109:E1848–E1857. doi: 10.1073/pnas.1200250109

14. Cyganek L, Tiburcy M, Sekeres K, Gerstenberg K, Bohnenberger H, Lenz C, Henze S, Stauske M, Salinas G, Zimmermann WH, et al. Deep phenotyping of human induced pluripotent stem cell–derived atrial and ventricular cardiomyocytes. JCI Insight. 2018;3:e99941. doi: 10.1172/jci.insight.99941

15. Reid CA, Boye SL, Hauswirth WW, Lipinski DM. miRNA-mediated post-transcriptional silencing of transgenes leads to increased adeno-associated viral vector yield and targeting specificity. Gene Ther. 2017;24:462–469. doi: 10.1038/gt.2017.50

16. Feyen DAM, McKeithan WL, Bruyneel AAN, Spiering S, Hörmann L, Ulmer B, Zhang H, Briganti F, Schweizer M, Hegyi B, et al. Metabolic maturation media improve physiological function of human iPSC-derived cardiomyocytes. Cell Rep. 2020;32. doi: 10.1016/j.celrep.2020.107925

17. Subramanian A, Tamayo P, Mootha VK, Mukherjee S, Ebert BL, Gillette MA, Paulovich A, Pomeroy SL, Golub TR, Lander ES, et al. Gene set enrichment analysis: a knowledge-based approach for interpreting genome-wide expression profiles. Proc Natl Acad Sci U S A. 2005;102:15545–15550. doi: 10.1073/pnas.0506580102

18. MSigDB 2023.1.Hs - GSEA-MSigDB Documentation.

19. Zhang L, Kulkarni P, Farshidfar F, Tingley W, Hoey T, Wang W, Priest JR, Figarska SM. Combining genetic proxies of drug targets and time-to-event analyses from longitudinal observational data to identify target patient populations. BMC Cardiovasc Disord. 2025;25:353. doi: 10.1186/s12872-025-04753-1

20. Foley CN, Staley JR, Breen PG, Sun BB, Kirk PDW, Burgess S, Howson JMM. A fast and efficient colocalization algorithm for identifying shared genetic risk factors across multiple traits. Nat Commun. 2021;12:764. doi: 10.1038/s41467-020-20885-8

21. Pirruccello JP, Di Achille P, Nauffal V, Nekoui M, Friedman SF, Klarqvist MDR, Chaffin MD, Weng LC, Cunningham JW, Khurshid S, et al. Genetic analysis of right heart structure and function in 40,000 people. Nat Genet. 2022;54:792–803. doi: 10.1038/s41588-022-01090-3

22. Tadros R, Francis C, Xu X, Vermeer AMC, Harper AR, Huurman R, Kelu Bisabu K, Walsh R, Hoorntje ET, te Rijdt WP, et al. Shared genetic pathways contribute to risk of hypertrophic and dilated cardiomyopathies with opposite directions of effect. Nat Genet. 2021;53:128–134. doi: 10.1038/s41588-020-00762-2

23. Evangelou E, Warren HR, Mosen-Ansorena D, Mifsud B, Pazoki R, Gao H, Ntritsos G, Dimou N, Cabrera CP, Karaman I, et al. Genetic analysis of over 1 million people identifies 535 new loci associated with blood pressure traits. Nat Genet. 2018;50:1412–1425. doi: 10.1038/s41588-018-0205-x

24. Zhou W, Kanai M, Wu KHH, Rasheed H, Tsuo K, Hirbo JB, Wang Y, Bhattacharya A, Zhao H, Namba S, et al. Global Biobank Meta-analysis Initiative: Powering genetic discovery across human disease. Cell Genomics. 2022;2:100192. doi: 10.1016/j.xgen.2022.100192

25. Lee JH, Protze SI, Laksman Z, Backx PH, Keller GM. Human pluripotent stem cell-derived atrial and ventricular cardiomyocytes develop from distinct mesoderm populations. Cell Stem Cell. 2017;21:179–194. doi: 10.1016/j.stem.2017.07.003

26. Christophersen IE, Rienstra M, Roselli C, Yin X, Geelhoed B, Barnard J, Lin H, Arking DE, Smith A V., Albert CM, et al. Large-scale analyses of common and rare variants identify 12 new loci associated with atrial fibrillation. Nat Genet. 2017;49:946–952. doi: 10.1038/ng.3843

27. Nielsen JB, Fritsche LG, Zhou W, Teslovich TM, Holmen OL, Gustafsson S, Gabrielsen ME, Schmidt EM, Beaumont R, Wolford BN, et al. Genome-wide study of atrial fibrillation identifies seven risk loci and highlights biological pathways and regulatory elements involved in cardiac development. Am J Hum Genet. 2018;102:103–115. doi: 10.1016/j.ajhg.2017.12.003

28. Ntalla I, Weng LC, Cartwright JH, Hall AW, Sveinbjornsson G, Tucker NR, Choi SH, Chaffin MD, Roselli C, Barnes MR, et al. Multi-ancestry GWAS of the electrocardiographic PR interval identifies 202 loci underlying cardiac conduction. Nat Commun. 2020;11:2542. doi: 10.1038/s41467-020-15706-x

29. Mandyam MC, Soliman EZ, Alonso A, Dewland TA, Heckbert SR, Vittinghoff E, Cummings SR, Ellinor PT, Chaitman BR, Stocke K, et al. The QT interval and risk of incident atrial fibrillation. Heart Rhythm. 2013;10:1562–1568. doi: 10.1016/j.hrthm.2013.07.023

30. Farinha JM, Gupta D, Lip GYH. Frequent premature atrial contractions as a signalling marker of atrial cardiomyopathy, incident atrial fibrillation, and stroke. Cardiovasc Res. 2023;119:429–439. doi: 10.1093/cvr/cvac054

31. Voigt N, Trausch A, Knaut M, Matschke K, Varró A, Van Wagoner DR, Nattel S, Ravens U, Dobrev D. Left-to-right atrial inward rectifier potassium current gradients in patients with paroxysmal versus chronic atrial fibrillation. Circ Arrhythm Electrophysiol. 2010;3:472–480. doi: 10.1161/CIRCEP.110.954636

32. Kang GJ, Xie A, Kim E, Dudley SC. miR-448 regulates potassium voltage-gated channel subfamily A member 4 (KCNA4) in ischemia and heart failure. Heart Rhythm. 2023;20:730–736. doi: 10.1016/j.hrthm.2023.01.021

33. Liu B, Lou Q, Smith H, Velez-Cortes F, Dillmann WH, Knollmann BC, Armoundas AA, Györke S. Conditional up-regulation of SERCA2a exacerbates RyR2-dependent ventricular and atrial arrhythmias. Int J Mol Sci. 2020;21:2535. doi: 10.3390/ijms21072535

34. Yan J, Bare DJ, Desantiago J, Zhao W, Mei Y, Chen Z, Ginsburg K, Solaro RJ, Wolska BM, Bers DM, et al. JNK2, a newly-identified SERCA2 enhancer, augments an arrhythmic [Ca^2+^]_SR_ leak-load relationship. Circ Res. 2021;128:455–470. doi: 10.1161/CIRCRESAHA.120.318409

35. Kostin S, Klein G, Szalay Z, Hein S, Bauer EP, Schaper J. Structural correlate of atrial fibrillation in human patients. Cardiovasc Res. 2002;54:361–379. doi: 10.1016/S0008-6363(02)00273-0

36. Bikou O, Thomas D, Trappe K, Lugenbiel P, Kelemen K, Koch M, Soucek R, Voss F, Becker R, Katus HA, et al. Connexin 43 gene therapy prevents persistent atrial fibrillation in a porcine model. Cardiovasc Res. 2011;92:218–225. doi: 10.1093/cvr/cvr209

37. Espinoza-Lewis RA, Yu L, He F, Liu H, Tang R, Shi J, Sun X, Martin JF, Wang D, Yang J, et al. Shox2 is essential for the differentiation of cardiac pacemaker cells by repressing Nkx2-5. Dev Biol. 2009;327:376–385. doi:10.3390/biom12060859

38. Li H, Tang Q, Yang T, Wang Z, Li D, Wang L, Li L, Chen Y, Huang H, Zhang Y, et al. Segregation of morphogenetic regulatory function of *Shox2* from its cell fate guardian role in sinoatrial node development. Commun Biol. 2024;7:385. doi:10.1038/s42003-024-06039-2

39. Puskaric S, Schmitteckert S, Mori AD, Glaser A, Schneider KU, Bruneau BG, Blaschke RJ, Steinbeisser H, Rappold G. Shox2 mediates Tbx5 activity by regulating Bmp4 in the pacemaker region of the developing heart. Hum Mol Genet. 2010;19:4625–4633. doi:10.1093/hmg/ddq393

40. Gore-Panter SR, Hsu J, Hanna P, Gillinov AM, Pettersson G, Newton DW, Moravec CS, Van Wagoner DR, Chung MK, Barnard J, et al. Atrial fibrillation associated chromosome 4q25 variants are not associated with PITX2c expression in human adult left atrial appendages. PLoS ONE. 2014;9:e86245. doi:10.1371/journal.pone.0086245

41. Liang D, Xue J, Geng L, Zhou L, Lv B, Zeng Q, Xiong K, Zhou H, Xie D, Zhang F, et al. Cellular and molecular landscape of mammalian sinoatrial node revealed by single-cell RNA sequencing. Nat Commun. 2021;12:287. doi:10.1038/s41467-020-20448-x

42. van Eif VWW, Stefanovic S, van Duijvenboden K, Bakker M, Wakker V, de Gier-de Vries C, Zaffran S, Verkerk AO, Boukens BJ, Christoffels VM. Transcriptome analysis of mouse and human sinoatrial node cells reveals a conserved genetic program. Development. 2019;146:dev173161. doi:10.1242/dev.173161

43. Brunello L, Slabaugh JL, Radwanski PB, Ho HT, Belevych AE, Lou Q, Chen H, Napolitano C, Lodola F, Priori SG, et al. Decreased RyR2 refractoriness determines myocardial synchronization of aberrant Ca^2+^ release in a genetic model of arrhythmia. Proc Natl Acad Sci U S A. 2013;110:10312–10317. doi: 10.1073/pnas.1300052110

44. Keidar N, Elul Y, Schuster A, Yaniv Y. Visualizing and quantifying irregular heart rate irregularities to identify atrial fibrillation events. Front Physiol. 2021;12:637680. doi: 10.3389/fphys.2021.637680

45. Liu J, Ye L, Li Q, Wu X, Wang B, Ouyang Y, Yuan Z, Li J, Lin C. Synaptopodin-2 suppresses metastasis of triple-negative breast cancer via inhibition of YAP/TAZ activity. J Pathol. 2018;244:71–83. doi: 10.1002/path.4995

46. Zheng M, Li RG, Song J, Zhao X, Tang L, Erhardt S, Chen W, Nguyen BH, Li X, Li M, et al. Hippo-Yap signaling maintains sinoatrial node homeostasis. Circulation. 2022;146:1694–1711. doi: 10.1161/CIRCULATIONAHA.121.058777

47. Kashihara T, Sadoshima J. Role of YAP/TAZ in energy metabolism in the heart. J Cardiovasc Pharmacol. 2019;74:483–490. doi: 10.1097/FJC.0000000000000736

48. Ikeda S, Mizushima W, Sciarretta S, Abdellatif M, Zhai P, Mukai R, Fefelova N, Oka SI, Nakamura M, Del Re DP, et al. Hippo deficiency leads to cardiac dysfunction accompanied by cardiomyocyte dedifferentiation during pressure overload. Circ Res. 2019;124:292–305. doi: 10.1161/CIRCRESAHA.118.314048

49. Liu L, Huang S, Du Y, Zhou H, Zhang K, He J. Lats2 deficiency protects the heart against myocardial infarction by reducing inflammation and inhibiting mitochondrial fission and STING/p65 signaling. Int J Biol Sci. 2023;19:3428–3440. doi:10.7150/ijbs.84426

50. Matsui Y, Nakano N, Shao D, Gao S, Luo W, Hong C, Zhai P, Holle E, Yu X, Yabuta N, et al. Lats2 is a negative regulator of myocyte size in the heart. Circ Res. 2008;103:1309–1318. doi: 10.1161/CIRCRESAHA.108.180042

51. Aguet F, Barbeira AN, Bonazzola R, Brown A, Castel SE, Jo B, Kasela S, Kim-Hellmuth S, Liang Y, Oliva M, et al. The GTEx Consortium atlas of genetic regulatory effects across human tissues. Science. 2020;369:1318–1330. doi: 10.1126/science.aaz1776

52. Schmitt J, Duray G, Gersh BJ, Hohnloser SH. Atrial fibrillation in acute myocardial infarction: A systematic review of the incidence, clinical features and prognostic implications. Eur Heart J. 2009;30:1038–1045. doi: 10.1093/eurheartj/ehn579

53. Holmes MV. Human genetics and drug development. N Engl J Med. 2019;380:1076–1079. doi: 10.1056/NEJMe1901565

54. O’Sullivan JW, Raghavan S, Marquez-Luna C, Luzum JA, Damrauer SM, Ashley EA, O’Donnell CJ, Willer CJ, Natarajan P. Polygenic risk scores for cardiovascular disease: a scientific statement from the American Heart Association. Circulation. 2022;146:E93–E118. doi: 10.1161/CIR.000000000000107

55. Shah S, Henry A, Roselli C, Lin H, Sveinbjörnsson G, Fatemifar G, Hedman ÅK, Wilk JB, Morley MP, Chaffin MD, et al. Genome-wide association and Mendelian randomisation analysis provide insights into the pathogenesis of heart failure. Nat Commun. 2020;11:163. doi:10.1038/s41467-019-13690-5

56. Yamada H, Osaka H, Tatsumi N, Araki M, Abe T, Kaihara K, Takahashi K, Takashima E, Uchihashi T, Naruse K, et al. Direct binding of synaptopodin 2-like protein to alpha-actinin contributes to actin bundle formation in cardiomyocytes. Cells. 2024;13:1373. doi:10.3390/cells13161373

