## Supplemental Information for "SYNPO2L Isoforms Regulate the Action Potential Characteristics and Contractility of Atrial Cardiomyocytes via YAP Signaling to Modulate Atrial Fibrillation Risk"

Supplemental Figures S1-S9

Supplemental Tables S1-S3

**Table S1.** Antibody supplier and dilution information.

| Primary Antibodies | | |
|---|---|---|
| **Antibody** | **Dilution** | **Supplier (Part #)** |
| SYNPO2LA | 1:100 | Genscript (Custom) |
| SYNPO2LB | 1:100 | Genscript (Custom) |
| LATS2 | 1:200 | Abcam (ab243657) |
| MLC2A/MYL7 | 1:500 | Invitrogen (PA530789) |
| MLC2V/MYL2 | 1:500 | Invitrogen (PA586045) |
| GAPDH | 1:2500 | Invitrogen (437000) |
| SYNPO2LA/B | 1:300 | Proteintech (501733440) |
| **Secondary Antibodies** | | |
| **Antibody** | **Dilution** | **Supplier (Part #)** |
| Goat Anti-Rabbit | 1:2500 | Licor (926-32211) |
| Goat Anti-Mouse | 1:2500 | Licor (926-32210) |

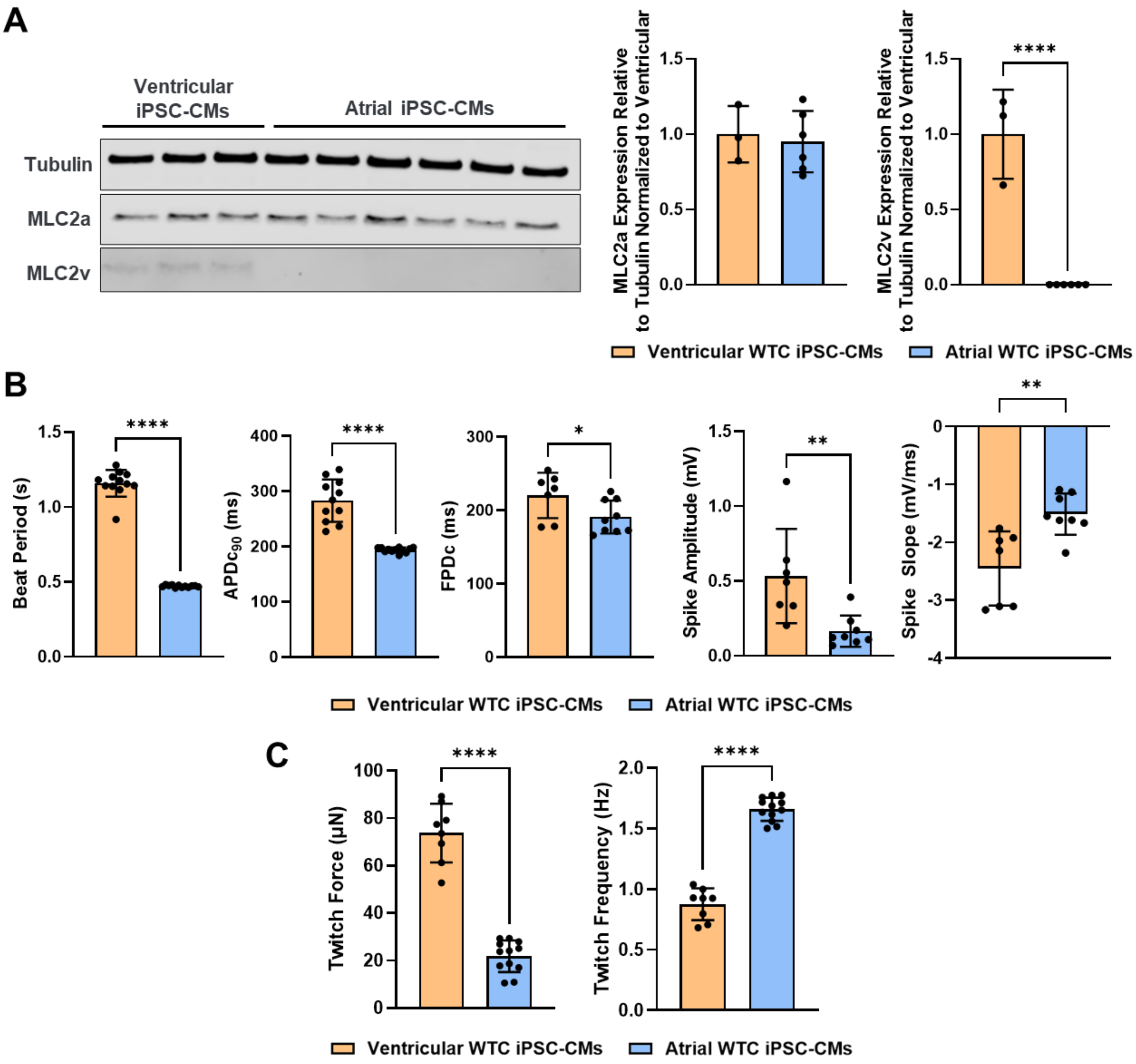


**Figure S1. Atrial hiPSC-CMs display molecular and functional markers of an atrial cardiomyocyte phenotype.** (A) Protein expression of MLC2v is significantly reduced in atrial lineage cardiomyocytes. (B) Atrial cardiomyocytes display reduced beat periods, action potential durations, field potential durations, spike amplitudes, and spike slopes relative to ventricular cardiomyocytes. (C) Twitch forces produced by atrial cardiomyocytes are also significantly reduced, and the increase in twitch frequency corresponds with the reduced beat periods observed. n≥3 biological replicates for western blot data; n≥7 biological replicates for MEA data; n≥8 biological replicates for EHT data. *p<0.05, **p<0.005, ****p<0.0001 as determined by Student's t-test.

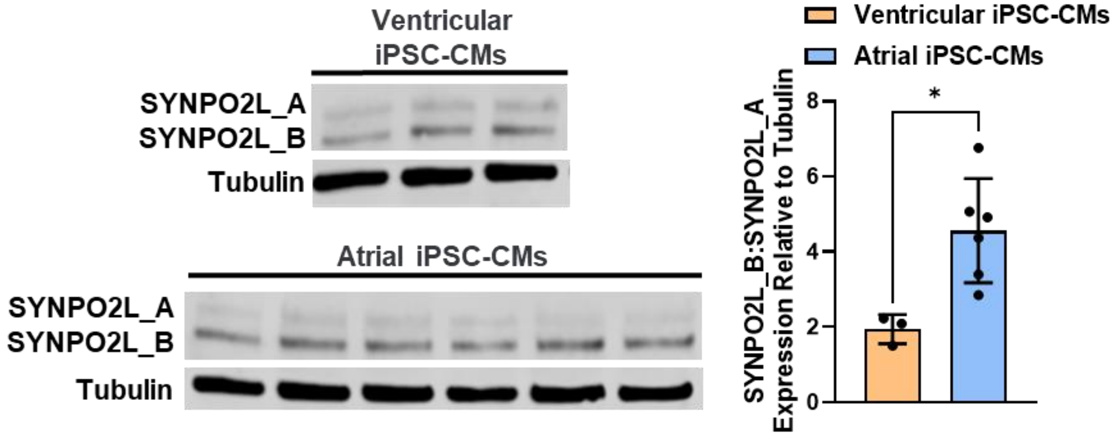


**Figure S2. Relative expression of SYNPO2L isoforms differs in atrial cardiomyocytes.** At the protein level, the ratio of SYNPO2L_B to SYNPO2L_A is greater in atrial hiPSC-CMs than in ventricular hiPSC-CMs. n=6 biological replicates. n≥3 biological replicates. *p<0.05 as determined by Student's t-test.

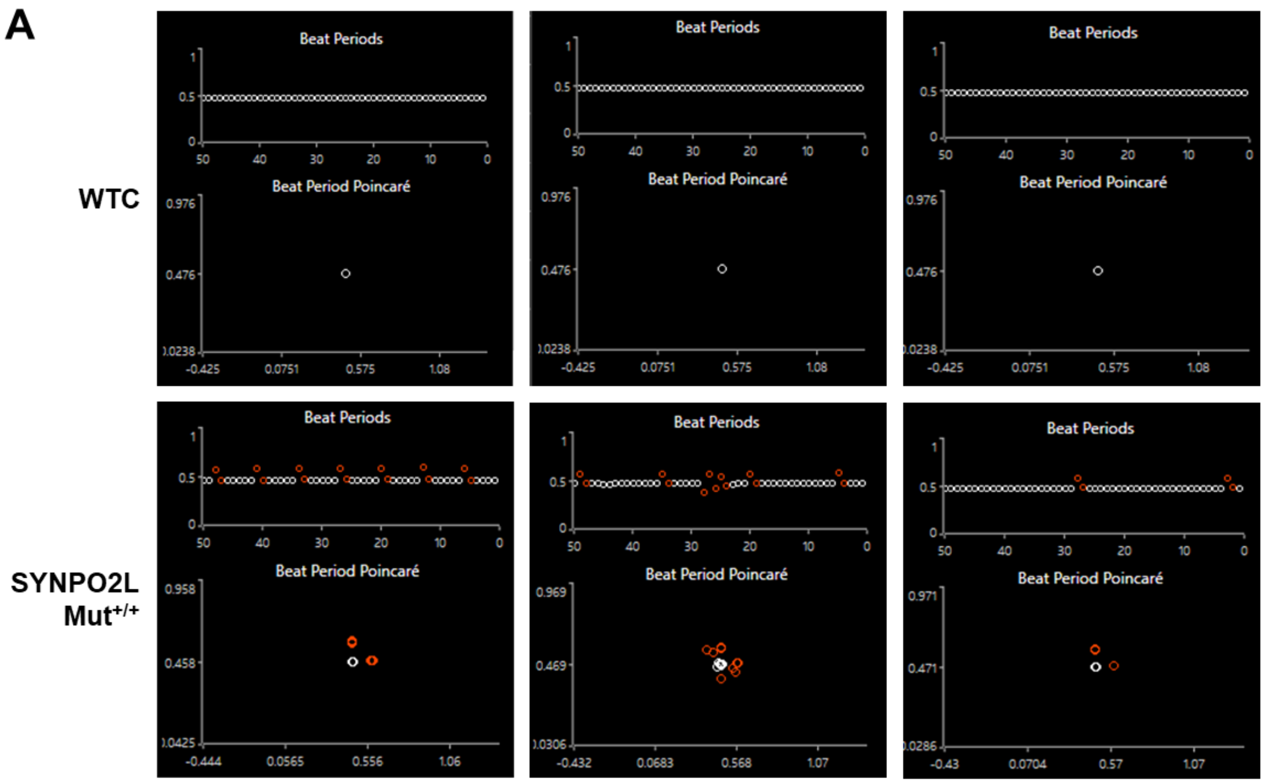


B

| Minutes | WTC | *SYNPO2L*$^{SS/SS}$ |
|---|---|---|
| 0 | 0/12 | 0/12 |
| 15 | 0/12 | 5/12 |
| 30 | 0/12 | 4/12 |
| 45 | 0/12 | 5/12 |
| 60 | 0/12 | 4/12 |
| 75 | 0/12 | 2/12 |
| 90 | 0/12 | 0/12 |

Number of Wells Producing Extra Beats/Total Number of Wells

**Figure S3. Atrial *SYNPO2L*$^{SS/SS}$ hiPSC-CMs beat more irregularly during long term pacing.** (A) Increased instances of beat period instability were observed in *SYNPO2L*$^{SS/SS}$ atrial cardiomyocyte monolayers as measured by MEA. (B) Over a 90-minute pacing period, SYNPO2L mutant atrial hiPSC-CMs were more prone to produce extra beats compared to no extra beats produced by the WTC control atrial hiPSC-CMs. p=0.007 as determined by a linear fit model accounting for time.

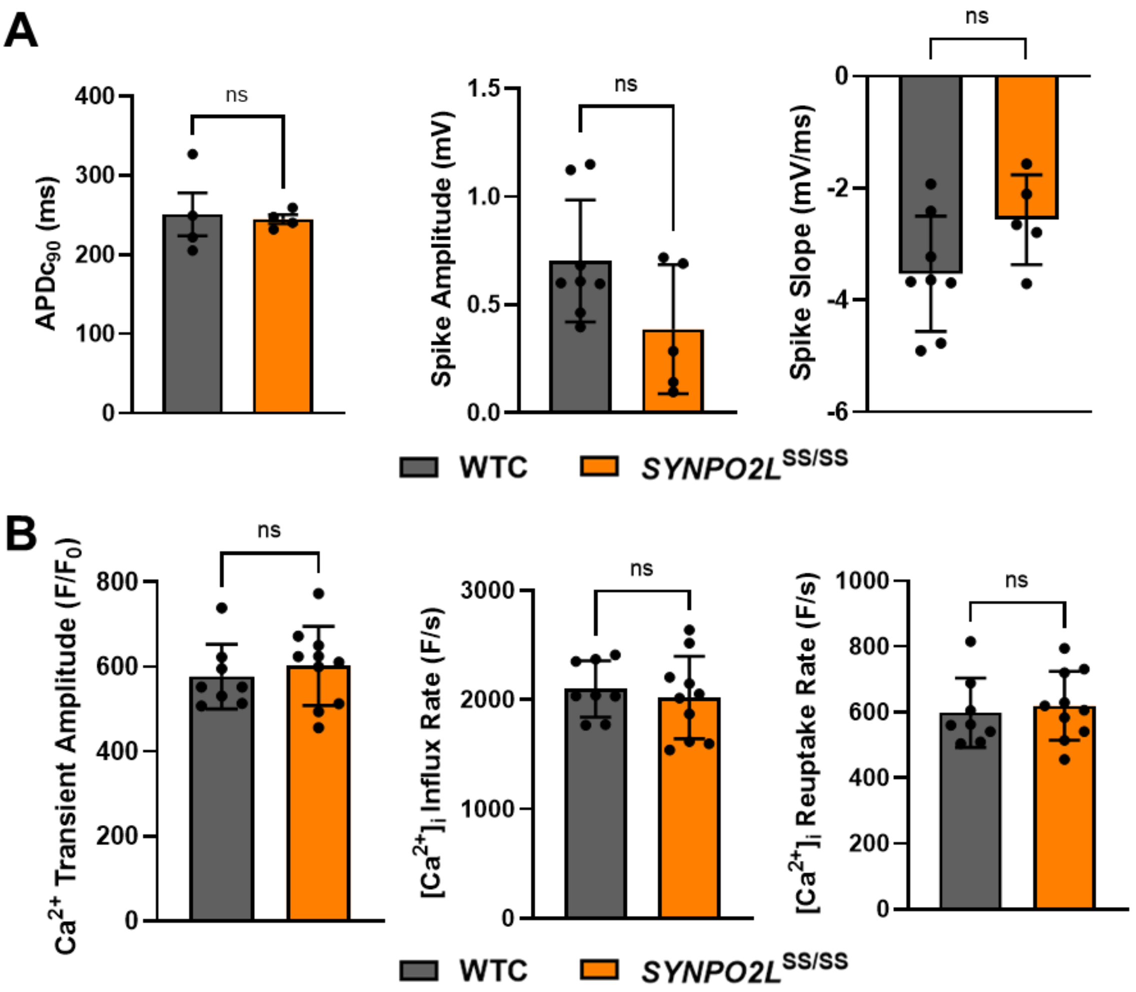


**Figure S4. Ventricular *SYNPO2L*$^{ss/ss}$ hiPSC-CMs display only slight differences in electrophysiological function compared to WTC.** (A) Measures of actional potential durations, spike amplitudes, and spike slopes show subtle, but not significant, changes, with spike amplitudes and spike slopes slightly reduced in mutant cells. (B) Similarly, calcium handling metrics show no difference between wild-type and mutant ventricular cardiomyocytes. n≥5 biological replicates for MEA data; n≥8 biological replicates for calcium imaging data.

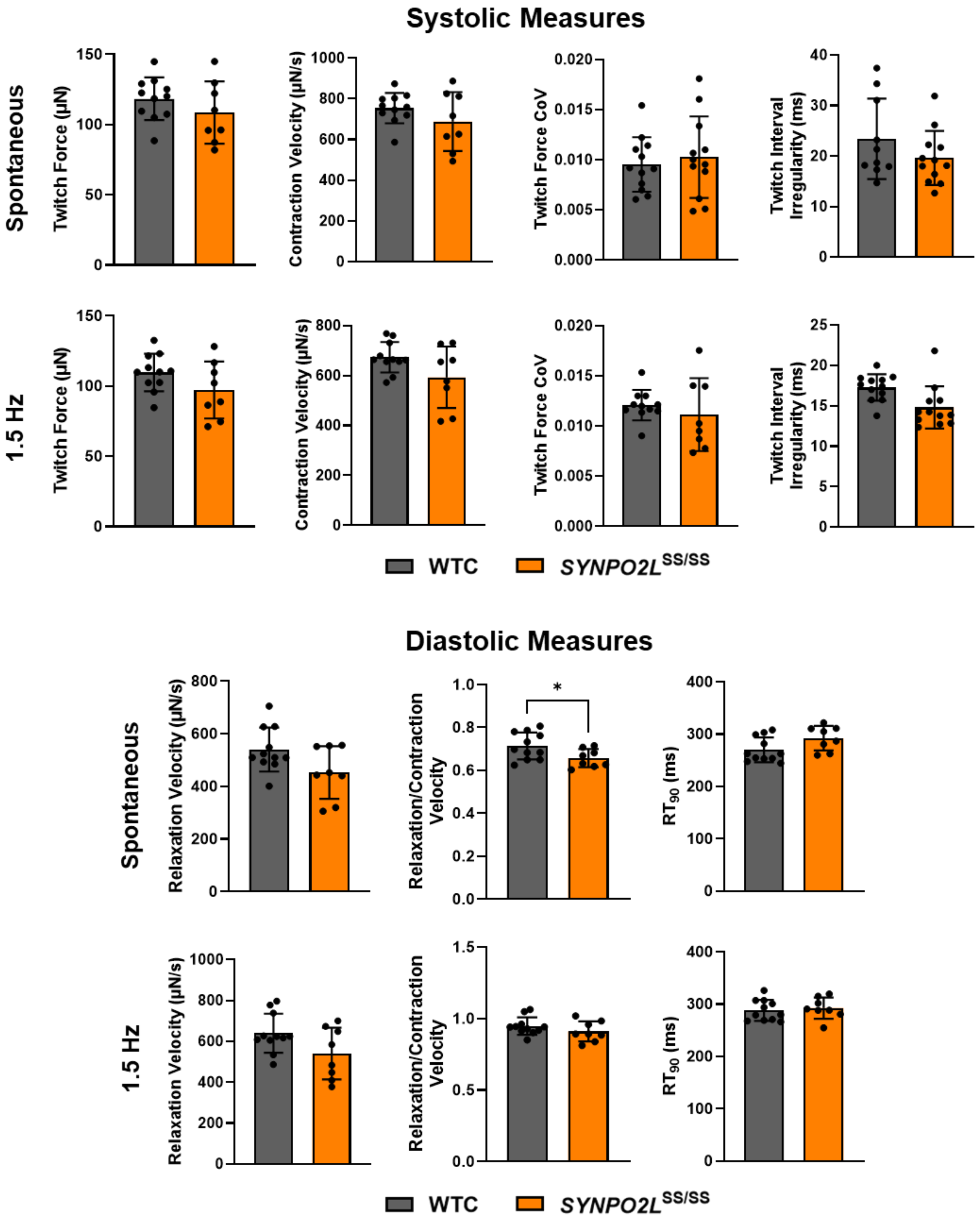


**Figure S5. Ventricular *SYNPO2L*$^{SS/SS}$ hiPSC-CMs display mild differences in diastolic function but unchanged systolic function.** Under both paced and un-paced conditions, EHTs produced from ventricular mutant hiPSC-CMs show no discernible differences in systolic contractile behavior compared to ventricular WTC EHTs. Similarly, differences in diastolic behavior are slight, with relaxation/contraction velocities the only metric with a significant decline in *SYNPO2L*$^{SS/SS}$ tissues. n≥8 biological replicates. *$p<0.05$ as determined by Student's t-test.

**Table S2.** Transcriptional response of downstream targets of YAP signaling in *SYNPO2L*$^{SS/SS}$ atrial iPSC-CMs.

| Gene | t-statistic | p.value (<0.05) | Mean WTC EHTs | Mean *SYNPO2L*$^{SS/SS}$ EHTs | Log2FC | Q |
|---|---|---|---|---|---|---|
| **ITGA9** | -19.26 | 0.00 | 3.34 | 2.54 | -0.80 | 0.00 |
| **ITGA5** | -12.03 | 0.00 | 6.22 | 5.12 | -1.10 | 0.00 |
| **CCN1** | -11.29 | 0.00 | 5.21 | 3.53 | -1.68 | 0.00 |
| **TGFB2** | -10.41 | 0.00 | 3.95 | 2.73 | -1.21 | 0.00 |
| **CCN2** | -7.63 | 0.00 | 5.70 | 3.72 | -1.98 | 0.00 |
| **KDR** | -7.37 | 0.00 | 2.57 | 1.26 | -1.30 | 0.00 |
| **CDH11** | -7.26 | 0.00 | 4.16 | 3.22 | -0.94 | 0.00 |
| **AMOT** | -7.16 | 0.00 | 1.32 | 0.84 | -0.48 | 0.00 |
| **KLF4** | -6.42 | 0.00 | 1.09 | 0.28 | -0.81 | 0.00 |
| **CDH2** | -6.14 | 0.00 | 7.54 | 7.23 | -0.31 | 0.00 |
| **ITGB5** | -4.97 | 0.00 | 4.92 | 4.53 | -0.39 | 0.01 |
| **ITGA1** | -4.79 | 0.00 | 1.59 | 1.01 | -0.58 | 0.01 |
| **MYOD1** | -4.79 | 0.00 | 0.91 | 0.41 | -0.50 | 0.01 |
| **ITGA2** | -4.48 | 0.00 | 0.34 | 0.16 | -0.18 | 0.01 |
| **CTNNA1** | -4.23 | 0.00 | 8.69 | 8.57 | -0.12 | 0.01 |
| **USP9X** | -3.89 | 0.00 | 5.12 | 4.92 | -0.19 | 0.02 |
| **ITGAX** | -3.67 | 0.01 | 0.56 | 0.09 | -0.47 | 0.04 |
| **CDH8** | -3.62 | 0.00 | 2.89 | 2.15 | -0.73 | 0.02 |
| **CDH19** | -3.46 | 0.02 | 0.07 | 0.00 | -0.07 | 0.07 |

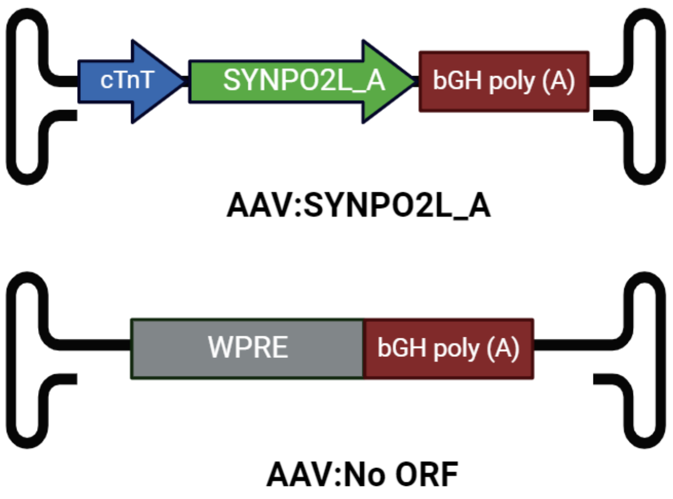


**Figure S6. Cassettes for AAV:SYNPO2L_A and AAV:No ORF.** Regulatory features of the AAV:SYNPO2L_A include a human cardiac troponin promoter and a bGH poly(A) tail. The AAV:No ORF cassette does not contain a promoter nor an open reading frame, but does contain a WPRE and bGH poly(A) tail.

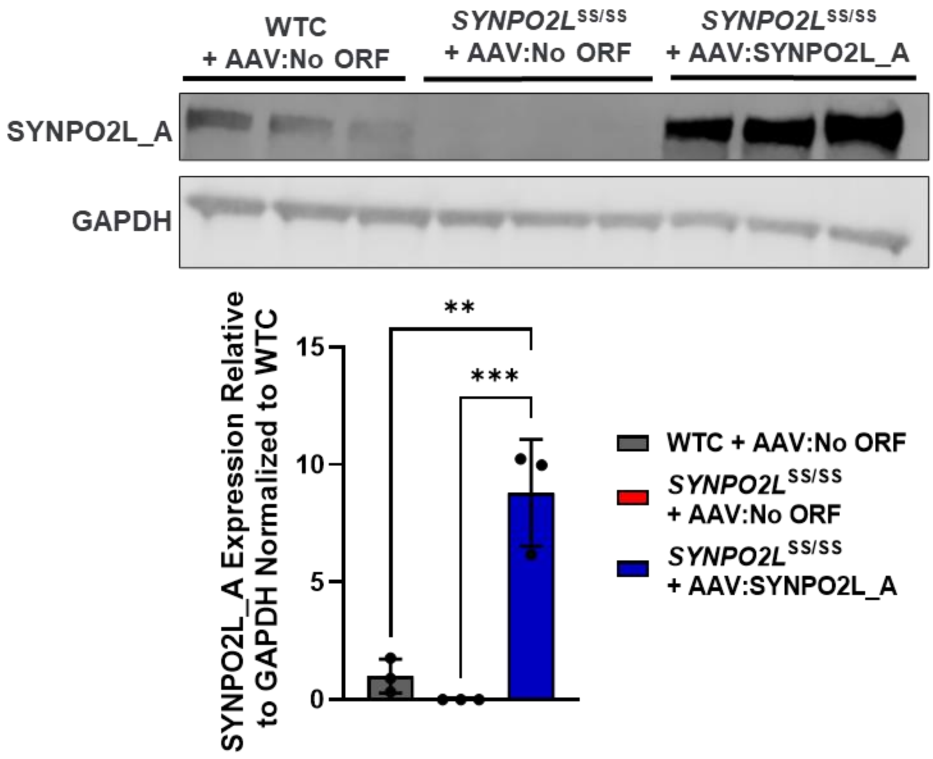


**Figure S7. Quantification of SYNPO2L_A Overexpression post-transduction with AAV.** Transduction with a 5K MOI of AAV:SYNPO2L_A in *SYNPO2L*$^{SS/SS}$ cardiomyocytes led to an approximately 9-fold increase in SYNPO2L_A expression relative to WTC. n=3 biological replicates. **p<0.005, ***p<0.001 as determined by one-way ANOVA.

| Minutes | *SYNPO2L*$^{SS/SS}$ + AAV:No ORF | *SYNPO2L*$^{SS/SS}$ + AAV:SYNPO2L_A |
|---|---|---|
| 0 | 5/8 | 4/8 |
| 15 | 5/8 | 1/8 |
| 30 | 4/8 | 3/8 |
| 45 | 2/8 | 2/8 |
| 60 | 3/8 | 2/8 |
| 75 | 3/8 | 1/8 |
| 90 | 4/8 | 2/8 |

Number of Wells Producing Extra Beats/Total Number of Wells

**Figure S8. Overexpression of SYNPO2L_A in mutant cardiomyocytes partially restores rhythmicity.** Treatment with AAV:SYNPO2L_A resulted in a 21.4% reduction in aberrant beats over a 90-minute pacing period. p=0.006 as determined by a linear fit model accounting for time.

**Table S3.** A splice QTL for SYNPO2L SNP rs60632610 co-localizes with a variety of related traits.

| Traits | Posterior Prob | Regional Prob | Candidate SNP | Posterior explained by SNP |
|---|---|---|---|---|
| LV Ejection Fraction, Dilated Cardiomyopathy, Atrial Fibrillation, Diastolic Blood Pressure, Systolic Blood Pressure, Heart Failure (European Population), short axis pulmonary artery diameter | 0.3166 | 0.5606 | **rs60632610** | 0.9793 |
